# Single-cell and spatial multi-omics mapping of human ovarian aging across young and menopausal transition

**DOI:** 10.64898/2026.11.12.765495

**Authors:** Jingjing Li, Cheng Wu, Yue Zhang, Jianing Wu, Huiyu Fan, Chunming Li, Cheng Chen, Yijun Zhou, Linjuan Ma, Jianzhong Su, Jianhong Zhou, Yunlong Ma, Yizhou Huang

**Author notes:** Equal contribution (Jingjing Li, Cheng Wu).

## Abstract

Ovarian aging is an early feature of human aging with broad consequences for women’s health, yet its cellular dynamics across the menopausal transition remain unclear. We integrated single-cell transcriptomic, chromatin accessibility and spatial transcriptomic profiles across young, menopausal-transition and postmenopausal human ovaries. Granulosa cells (GCs) exhibited the most pronounced age-associated transcriptional remodelling. The menopausal transition was characterized by reduced DNA-repair activity and elevated stress-response programs in GCs, alongside broader changes in hormone-response and steroid-metabolic programs during ovarian aging. Integrated regulatory analyses implicated *HEYL*, a GC-enriched transcription factor, with reduced accessibility at a putative regulatory element during aging. Genetic analyses associated *HEYL* and its predicted upstream regulators with age at natural menopause, while spatial mapping revealed enrichment of menopause-associated genetic signals in GC-rich regions. *HEYL* depletion increased senescence-associated phenotypes in human granulosa-like cells and primary mouse GCs, whereas overexpression attenuated oxidative-stress-induced senescence. Perturbation of its orthologue *Hey* produced consistent effects in *Drosophila* ovaries. Together, our atlas reveals stage-associated granulosa-cell remodeling and implicates *HEYL* in ovarian aging.

## Introduction

The ovary is among the earliest human organs to exhibit age-associated functional decline, characterized by a progressive depletion of the follicular reserve and deterioration in oocyte quality ^1^. This decline accelerates with advancing reproductive age and ultimately culminates in menopause ^1^. Beyond its effects on fertility, ovarian aging has broader implications for women’s health. Menopause-associated hormonal changes have been linked to alterations in skeletal, cardiovascular, metabolic and neurological health ^2^. Understanding the cellular and molecular mechanisms underlying ovarian aging is therefore important for elucidating the relationship between reproductive aging and women’s health across the lifespan.

Ovarian aging involves multiple hallmarks of aging, including genomic instability, cellular senescence, mitochondrial dysfunction, epigenetic alterations, and disrupted intercellular communication^3–5^. Genome-wide association studies (GWAS) of age at natural menopause (ANM) have identified 290 loci, implicating DNA damage response pathways in reproductive aging ^6^. However, how these aging-related processes and genetic mechanisms operate across distinct ovarian cell populations remains incompletely understood. Single-cell and single-nucleus transcriptomics, complemented by spatial transcriptomics and chromatin accessibility profiling, have begun to characterize cell-type-specific molecular changes during ovarian aging ^7–11^,. Integrated transcriptomic and chromatin accessibility analyses have implicated mTOR as a prominent ovarian aging pathway and increased activity of the transcription factor *CEBPD* in the aged ovaries ^11^. Complementary single-cell and spatiotemporal transcriptomic studies have identified *FOXP1* as a protective regulator of ovarian aging that declines with age and suppresses cellular senescence through transcriptional repression of *CDKN1A*^12^.

Despite these advances, the cellular and regulatory changes associated with the menopausal transition remain insufficiently understood. Previous single-cell studies have primarily investigated ovarian aging using broad chronological age groups, with women aged 42-54 years ^13^ or 49-54 years ^11^ grouped into older cohorts without explicitly distinguishing those undergoing the menopausal transition from those who were already postmenopausal. Such grouping may obscure stage-associated molecular changes, leaving unclear how ovarian cell populations and their transcriptional and regulatory programs are remodelled during the menopausal transition. Computational approaches ^14–19^ integrating GWAS summary statistics with single-cell or spatial transcriptomic data have enabled the identification of trait-associated cell types and spatial regions. However, the link between menopause-associated genetic signals and stage-specific regulatory programs within the spatial context of the human ovary remains largely unresolved. Integrating transcriptomic, chromatin accessibility and spatial profiles with ANM genetic associations, together with functional investigation of prioritized regulators could further insights into the mechanisms underlying ovarian aging.

Here, we present an integrated single-cell and spatial multi-omic atlas of human ovarian aging across young, menopausal-transition and postmenopausal stages. By combining single-cell transcriptomics, chromatin accessibility profiling and spatial transcriptomics, we characterized stage-associated cellular and regulatory changes, identifying granulosa cells (GCs) as a major site of remodelling during the menopausal transition. Integration of chromatin accessibility, regulatory networks and reproductive aging genetics further implicated *HEYL* as a candidate regulator of GC-associated aging programs. Functional perturbation in human granulosa-like cells, primary mouse GCs and *Drosophila* ovaries supported a role for *HEYL/Hey* in modulating senescence-associated phenotypes. Together, our study establishes a stage-resolved molecular atlas of human ovarian aging, providing insights into the cellular and regulatory changes associated with the menopausal transition.

## Results

### Single-cell multi-omics atlas of the human ovary across the menopausal transition

We generated a single-cell and spatial multi-omics atlas of human ovarian aging across three reproductive stages: young, menopausal transition (transition) and postmenopausal (post) (**Fig. 1a**). We performed single-cell RNA sequencing (scRNA-seq) and single-cell ATAC sequencing (scATAC-seq) on 15 ovarian samples each (five per group), together with Stereo-seq spatial transcriptomics on one tissue section per group (**Supplementary Table 1**). Donors were stratified according to age, menstrual history and serum follicle-stimulating hormone (FSH) levels, with the transition group comprising non-menopausal women in the late reproductive-aging period. Serum FSH levels increased across the three groups and positively correlated with age, supporting the endocrine stratification of the cohort (**Supplementary Fig. 1**).

**Figure 1:**
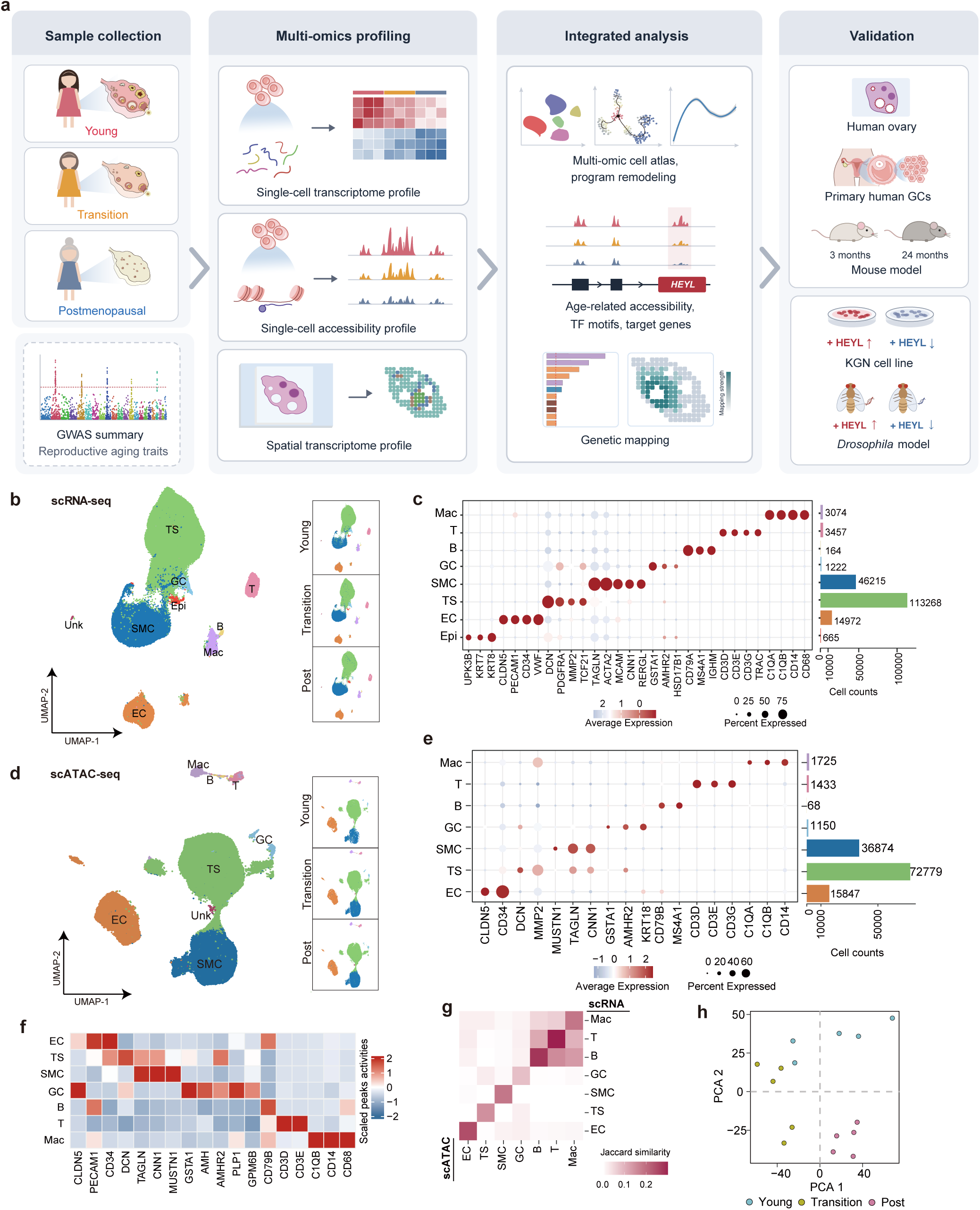
The single-cell atlas of the human ovary across the menopausal transition. **(a)** Schematic overview of the study design. **(b)** Uniform manifold approximation and projection (UMAP) of scRNA-seq cells, colored by annotated cell type (left) and split by age group (right). **(c)** Dot plot showing canonical marker gene expression across cell types (color indicates scaled mean expression; dot size indicates percentage of cells expressing each gene). Bar plot indicates the number of cells per cell type. **(d)** UMAP of scATAC-seq cells colored by cell type (left) and split by age group (right). **(e)** Dot plot of gene activity scores derived from scATAC-seq across cell types, with corresponding cell-type counts. **(f)** Correspondence between scRNA-seq- and scATAC-seq-derived cell-type annotations. **(g)** Principal component analysis (PCA) of pseudobulk scRNA-seq profiles. Samples colored by age group. **(h)** Scaled chromatin accessibility at marker-gene promoters (±0.5 kb of transcription start sites) across cell type. TS: theca/stromal cells, SMC: smooth muscle cells, EC: endothelial cells, GC: granulosa cells, Mac: macrophages, T: T cells, B: B cells, Epi: epithelial cells.

After quality control, we retained 183,394 cells from scRNA-seq and 129,936 cells from scATAC-seq (**Supplementary Fig. 2a,b**). Both modalities resolved major ovarian cell populations, including theca/stromal (TS), smooth muscle (SMC), endothelial (EC), granulosa (GC), macrophage (Mac), T, B, and epithelial (Epi) cells, annotated using canonical markers and gene activity scores (**Fig. 1b-e; Supplementary Table 2**). These cell-type annotations were consistent with previously reported human ovary atlases ^7,8,10^. TS and SMC represented the largest populations in both modalities (**Supplementary Fig. 2c,d**).

To assess concordance between the two modalities, we compared cell-type-specific transcriptional signatures with their corresponding chromatin accessibility profiles. Promoter regions of marker genes identified by scRNA-seq showed higher chromatin accessibility in the matching scATAC-seq cell types (**Fig. 1f**). Consistently, jaccard similarity analysis revealed greater overlap between cell-type-specific differentially expressed genes from scRNA-seq and differentially active genes from scATAC-seq in corresponding cell types (**Fig. 1g**). Principal-component analysis of cell-type pseudobulk gene expression profiles further showed that donors from the same reproductive stage tended to cluster together(**Fig. 1h**).

### Granulosa cells exhibit pronounced transcriptional remodelling during ovarian aging

We first examined changes in ovarian cell-type composition and gene expression across reproductive aging stages. Compositional analysis ^20^ revealed an increased relative abundance of SMCs and a decreased proportion of GCs with advancing age (**Fig. 2a,b**). These changes are consistent with the age-related depletion of ovarian follicles and previously reported stromal remodelling during ovarian aging ^2,8,21^.

**Figure 2:**
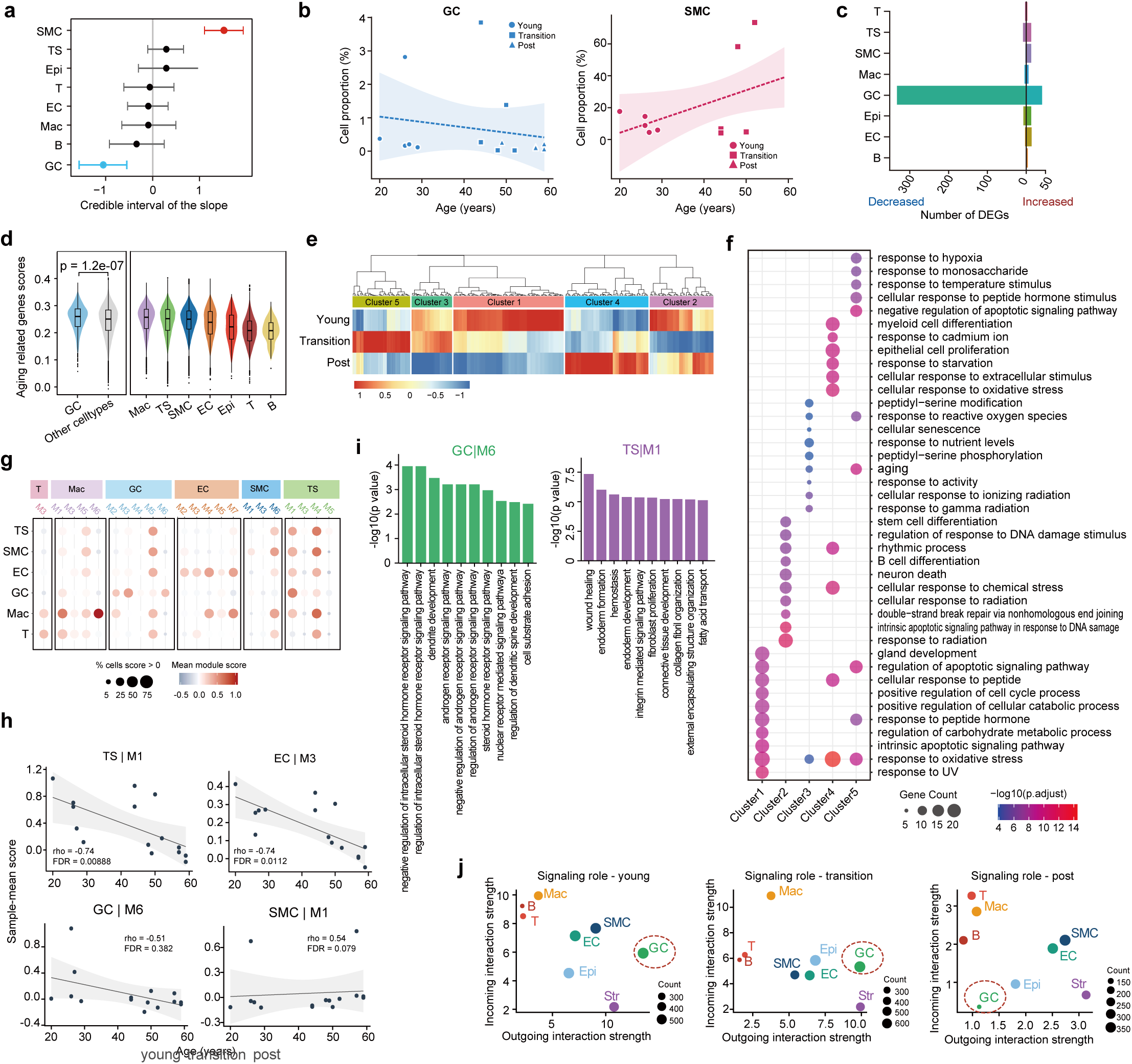
Altered cellular composition and transcriptional landscape of the ovary with age. **(a)** *sccomp* compositional analysis (post-menopausal versus young). Point represents posterior estimate of the age slope; bar represents credible interval. **(b)** Cell proportion versus age for GC and SMC cells (young/transition/post; regression line and 95% CI). **(c)** Number of age-associated DEGs per cell type (NEBULA, regression on continuous age; decreased, blue / increased, red). **(d)** Aging-related gene score (GenAge) by cell type. **(e)** Heatmap of aging-gene pseudobulk expression across young, transition, and post groups, with genes grouped into five clusters identified by unsupervised clustering. **(f)** GO enrichment of the five gene clusters in panel e. **(g)** hdWGCNA module specificity across cell types. Color means module scores. Size means percent of cells expressed. **(h)** Sample-mean activity versus age for four representative modules. Full module-age correlation in Supplementary Fig. 4c. **(i)** GO enrichment of two representative modules: the GC steroid-hormone-receptor module (GC|M6) and the stromal wound-healing/collagen module (TS|M1). **(j)** CellChat signaling-role scatter (incoming versus outgoing interaction strength) for young, transition and post-menopausal ovaries.

We next investigated age-associated transcriptional changes across ovarian cell types. Using NEBULA ^22^, a negative-binomial mixed-effects model, we identified the largest number of age-associated differentially expressed genes (DEGs) in GCs, despite their relatively low abundance (**Fig. 2c; Supplementary Table 3,4**). GCs also exhibited higher expression scores for aging-related genes curated from GenAge ^23^ compared with other ovarian cell types (*p* = 1.2 × 10^-7^; **Fig. 2d**). Functional enrichment analysis of age-associated DEGs in GCs identified pathways related to translation, ribosome biogenesis and DNA repair (**Supplementary Fig. 3a,b**), These findings identify GCs as a major site of transcriptional remodelling during ovarian aging.

To further characterize stage-associated transcriptional changes in GCs, we clustered GenAge genes based on their average expression across the three reproductive stages, identifying five distinct expression patterns (**Fig. 2e; Supplementary Table 5**). Cluster 1, enriched for cell proliferation and developmental processes, showed a progressive decrease in expression from young to post donors. Cluster 2, associated with DNA damage response and repair, exhibited lower expression in the transition group than either young and post group. Cluster 3, enriched for cellular senescence and reactive oxygen species responses, showed reduced expression primarily in the post group. Cluster 4, associated with oxidative stress and apoptotic signaling, showed progressively higher expression across reproductive stages. Cluster 5, enriched for hypoxia and cellular stress responses, showed the highest expression in the transition group (**Fig. 2f; Supplementary Fig. 3c**). These findings reveal distinct stage-associated transcriptional programs in GCs, including altered DNA damage-response and cellular stress-response programs during the menopausal transition.

To investigate coordinated transcriptional changes across ovarian cell types, we constructed cell-type-specific co-expression networks using hdWGCNA ^24^, identifying modules with distinct cell-type associations and biological functions (**Fig. 2g; Supplementary Fig. 4a,b**). Several modules associated with steroid hormone signalling, extracellular matrix organization and tissue remodelling exhibited age-related changes (**Supplementary Fig. 4c**). In GCs, a module enriched for steroid-hormone-receptor module (GC|M6) was negatively correlated with age. Similarly, a wound-healing-associated module in theca/stromal cells (TS|M1) and an extracellular matrix and collagen-associated module in endothelial cells (EC|M3) showed negative correlations with donor age. By contrast, a SMC module associated with muscle differentiation and contraction (SMC|M1) was positively associated with age (**Fig. 2i,h; Supplementary Fig. 4d**). These findings highlight distinct age-associated co-expression programs across ovarian cell types.

Finally, we examined changes in inferred cell-cell communication across reproductive stages. GCs exhibited lower incoming and outgoing interaction strengths in the post group than in the young and transition groups (**Fig. 2j**). At the tissue level, the postmenopausal group also showed fewer inferred interactions and lower overall interaction strength (**Supplementary Fig. 4h**). Pathway-level analyses revealed changes in several signaling networks, including collagen, laminin, MHC-I and APP signaling (**Supplementary Fig. 4e-g**). These analyses suggest altered intercellular communication during ovarian aging, with particularly reduced GC-associated signalling after menopause. Together, these findings highlight extensive remodelling of GCs during ovarian aging, encompassing changes in cellular abundance, transcriptional programs and intercellular communication, with distinct molecular alterations emerging during the menopausal transition.

### Granulosa-cell states and transcriptional programs during ovarian aging

Further subclustering of GCs identified four distinct populations: pre-granulosa cells (pGCs), cumulus GCs, mural GCs and atretic GCs (**Fig. 3a-c**). These subpopulations were characterized by established cell-type markers and distinct transcriptional profiles (**Fig. 3b**). Notably, atretic GCs exhibited expression of *IGFBP5*, *NR4A1* and *CDKN1A*, together with reduced expression of *FSHR* and *CYP19A1*, consistent with previously described atretic GC states ^25–27^. Pseudotime analysis positioned pGCs near the root of the inferred trajectory, with cumulus, mural and atretic GCs distributed along distinct branches (**Fig. 3d,e**). CytoTRACE ^28^ analysis revealed higher predicted differentiation potential in pGCs and lower scores in atretic GCs (**Fig. 3f**). The proportion of pGCs was higher in young ovaries, whereas atretic GCs became increasingly abundant in the transition and postmenopausal groups (**Fig. 3c**) Moreover, GCs from postmenopausal donors were more frequently distributed toward later pseudotime regions (**Fig. 3g,h**), suggesting age-associated shifts in GC states.

**Figure 3:**
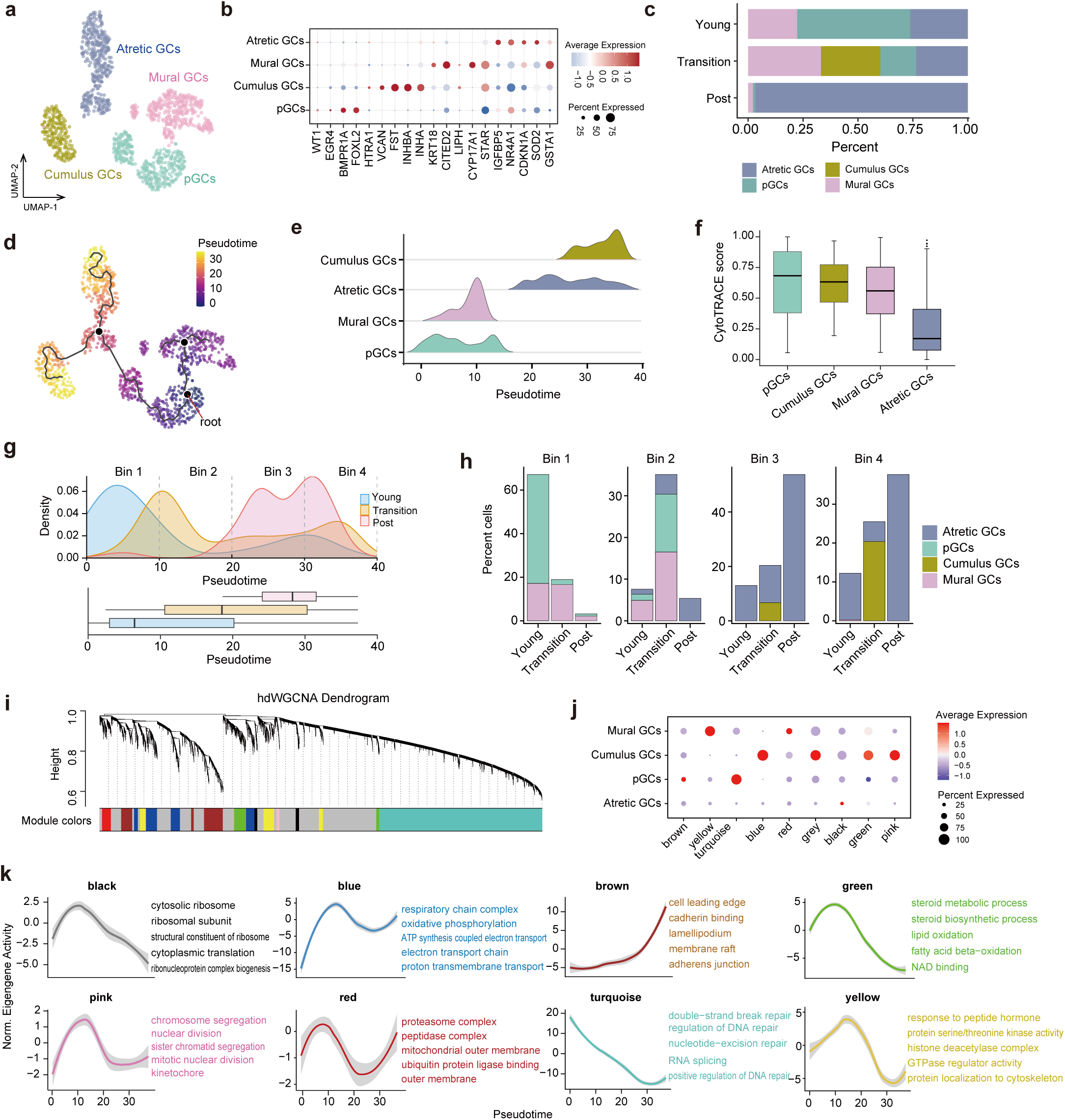
GC heterogeneity, trajectory and module dynamics during ovarian aging. **(a)** UMAP of GC subtypes: pre-granulosa cells (pGCs), cumulus, mural and atretic granulosa cells. **(b)** Dot plot of subtype marker genes. **(c)** GC subtype composition per age group. **(d)** Monocle3 pseudotime on the GC UMAP. **(e)** Pseudotime density per subtype. **(f)** CytoTRACE score per subtype. **(g)** Top, pseudotime density per age group divided into four equal-width bins; bottom, pseudotime distribution per age group (box plot). **(h)** GC subtype composition within each pseudotime bin across age groups. **(i)** hdWGCNA dendrogram and module colors for the GC co-expression network. **(j)** Module activity across GC subtypes (color, average expression; size, percent expressing). **(k)** Normalized module eigengene activity along pseudotime for each module, with representative GO terms labelled.

Next, GC-specific co-expression network analysis identified transcriptional modules with distinct activity patterns across GC subtypes and along the inferred pseudotime trajectory (**Fig. 3i-k; Supplementary Fig. 5**). Modules enriched for DNA repair and RNA splicing (turquoise), steroid metabolism and lipid oxidation (green), and peptide hormone responses (yellow) showed progressively lower activity along pseudotime. By contrast, modules associated with cell division (pink), ribosome biogenesis (black), oxidative phosphorylation (blue) and proteasome-related processes (red) exhibited non-monotonic patterns, with higher activity at early or intermediate pseudotime positions (**Fig. 3k**). Together, these findings reveal coordinated changes in transcriptional programs related to genome maintenance, metabolism, hormone responses and cellular homeostasis across GC states during ovarian aging.

### Cell-type-specific chromatin remodeling implicates *HEYL* in ovarian aging

Chromatin accessibility analysis identified widespread ovarian aging-associated differential accessible regions (DARs) across ovarian cell types (**Supplementary Fig. 6a; Supplementary Table 6**). Most DARs were located outside promoter regions, predominantly within intronic and intergenic regions (**Fig. 4a**). Motif enrichment analysis revealed distinct age-associated regulatory signatures inGCs. Regions displaying reduced accessibility with age were enriched for nuclear receptor motifs, including *NR5A1*, *ESRRA* and *ESRRB* (**Fig. 4b; Supplementary Fig. 6b,c**). *NR5A1* is a well-established regulator of ovarian steroidogenesis, whereas *ESRRA* and *ESRRB* have been implicated in mitochondrial and metabolic programs supporting steroid hormone production ^29–31^. In contrast, regions gaining accessibility were enriched for AP-1 family motifs (**Fig. 4c; Supplementary Fig. 6b,c**), including *FOS* and *JUN*, which are associated with cellular stress responses ^32,33^ .

**Figure 4:**
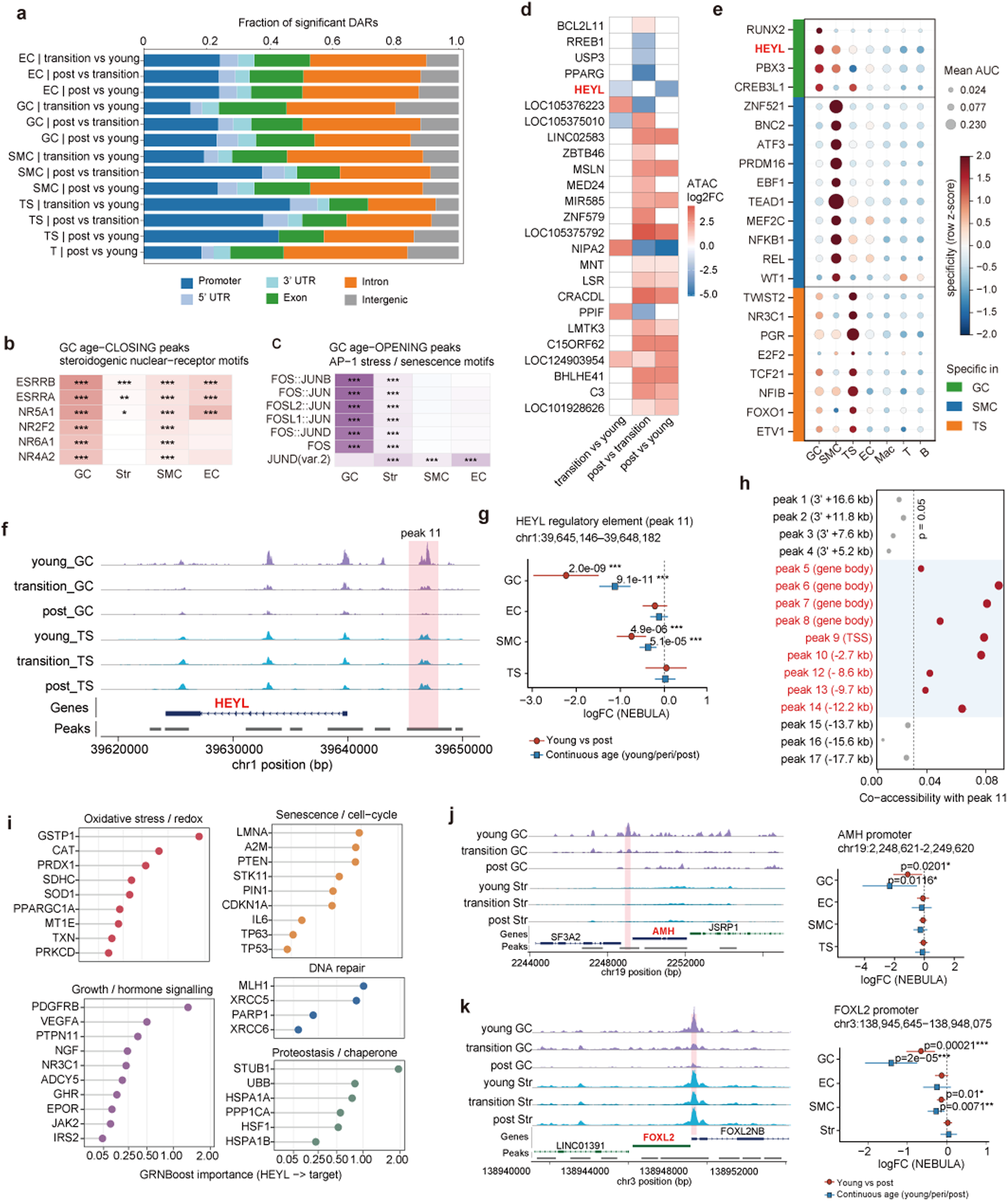
Cell-type-resolved chromatin remodeling converges on a granulosa *HEYL* program. **(a)** Genomic distribution of significant age-associated differentially accessible regions (DARs) in each cell type and contrast. DARs were identified with a logistic-regression test (Seurat FindMarkers; Bonferroni-adjusted *p* < 0.05 and |log2FC| > 0.1) and annotated relative to the nearest gene with ChIPseeker (promoter, ±3 kb of the TSS; downstream regions are included in intergenic). Bars show the fraction of DARs in each genomic category. Groups with fewer than 50 DARs were removed. **(b-c)** Cell-type-resolved TF motif enrichment in age-DARs (uniform Fisher vs each cell type’s peak background, JASPAR2020 CORE motifs). (b) Steroidogenic nuclear-receptor motifs (*ESRRB*, *ESRRA*, *NR5A1*, *NR2F2*, *NR6A1*, *NR4A2*) enriched among peaks that close with age; (c) AP-1/bZIP stress motifs enriched among peaks that open with age. Representative motifs are shown. **(d)** ATAC log2(fold-change) heatmap of top-25 GC DAR genes across age comparisons. **(e)** SCENIC+ eRegulon specificity × activity across cell types. **(f)** ATAC coverage across the *HEYL* locus in GC and TS cells, −5.5 kb element (peak 11) shaded. **(g)** Forest plot of peak 11 log2(fold change) with age in GC/EC/SMC/TS. **(h)** Single-cell co-accessibility of peak 11 with *HEYL*-locus peaks vs 2,000 random peaks (red, empirical *p* < 0.05). **(i)** Faceted lollipop of *HEYL* targets that are GenAge genes, by GRNBoost importance and functional category. **(j-k)** ATAC coverage (left) and forest plots (right) for the *AMH* **(j)** and *FOXL2* **(k)** promoters.

To further characterize age-associated changes in transcriptional regulation, we used chromVAR ^34^ to assess TF motif-associated chromatin accessibility across ovarian cell types. Motif activity scores showed predominantly negative associations with age across ovarian cell types (**Supplementary Fig. 6d**). In GCs, motifs associated with steroidogenic and lineage-related TFs, including *NR5A1*, *ESRRA*/*ESRRB* and FOX-family members, exhibited reduced accessibility with age, accompanied by decreased expression of the corresponding TF genes (**Supplementary Fig. 6e**). Moreover, age-associated changes in TF motif accessibility were broadly concordant with expression changes in their predicted target genes across ovarian cell types (**Supplementary Fig. 6f**). These findings suggest coordinated age-associated remodelling of TF-associated chromatin accessibility and transcriptional programs in the ovary.

To identify candidate regulators of GC aging, we integrated chromatin accessibility and gene expression using SCENIC+ ^35^. Among genes associated with age-related DARs in GCs, *HEYL* emerged as a candidate transcriptional regulator, with its inferred regulon exhibiting one of the highest GC-specificity scores across ovarian cell types (**Fig. 4d,e**). Consistently, HEY-family motif activity declined across reproductive aging stages in GCs, as assessed by ChromVAR (**Supplementary Fig. 7a**). At the *HEYL* locus, we identified several age-associated DARs, including a distal region approximately 5.5 kb from *HEYL* (peak 11; chr1:39,645,146-39,648,182) (**Fig. 4f,g; Supplementary Fig. 7b**). This region displayed reduced accessibility with age in GCs and showed co-accessibility with neighboring regions at the *HEYL* locus (**Fig. 4h**). Age-associated accessibility changes were also observed near established GC-associated genes, including *AMH* and *FOXL2* (**Fig. 4j,k**).

We next investigated the functional characteristics of predicted *HEYL* target genes. These targets were enriched for curated aging-related genes (**Fig. 4i**). Functional enrichment analysis implicated pathways related to mitochondrial oxidative phosphorylation and reactive oxygen species metabolism, as well as processes involved in cellular senescence, proteostasis and genome maintenance (**Supplementary Fig. 7c**). Together, these findings implicate *HEYL* as a candidate regulator of GC aging, linking age-associated chromatin remodelling to transcriptional programs involved in cellular maintenance and stress responses.

### Integration of reproductive aging genetics with ovarian regulatory programs

We integrated ovarian cell-type-specific transcriptional and chromatin accessibility changes with GWAS of reproductive aging and related traits, including age at natural menopause (ANM) ^6^, age at menarche ^36^, anti-Müllerian hormone levels ^37^, endometriosis ^38^, polycystic ovary syndrome ^39^, ovarian cancer ^40^ and breast cancer ^41^ (**Supplementary Table 7**). We observed enrichment of GWAS-associated genes among age-associated differentially expressed genes and genes linked to differentially accessible regions across multiple ovarian cell types (**Fig. 5a; Supplementary Fig. 8a,b**). This enrichment was particularly prominent for ANM and age at menarche in GC and TS gene sets.

**Figure 5:**
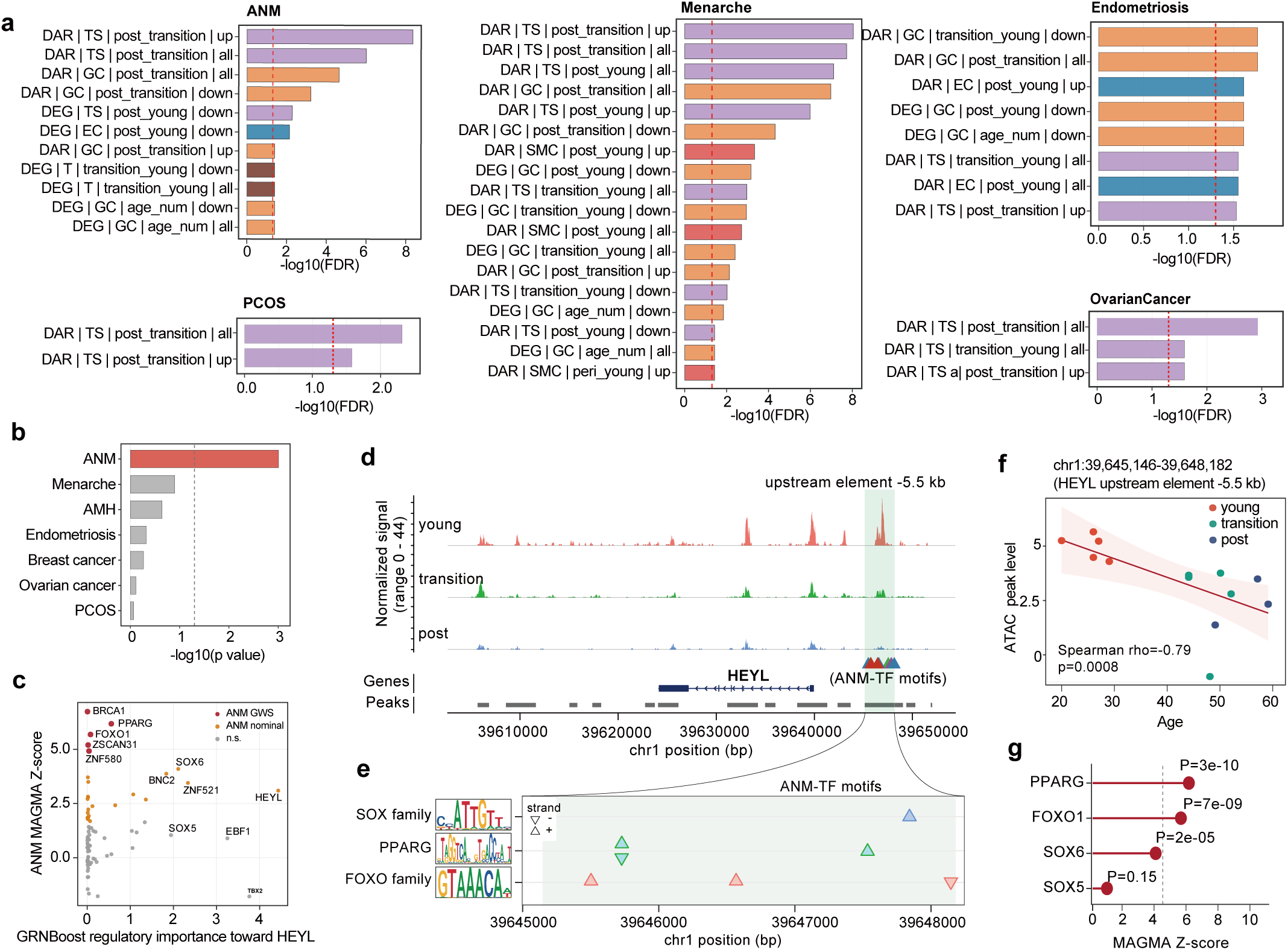
Ovarian aging genetic risk converges on the GC transcription factor *HEYL*. **(a)** Enrichment of ovarian trait genes in age-associated DAR/DEG gene sets across cell types, age contrasts and directions (one-sided Fisher’s exact test against per-cell type expressed-gene background; BH-FDR per trait; dashed line, FDR = 0.05). **(b)** *HEYL* gene-level association (MAGMA −log10(*p*-value)) across ovarian traits. **(c)** GRNBoost regulatory importance toward *HEYL* versus *ANM* gene-level MAGMA Z for 81 upstream transcription factors with *HEYL* as a predicted target in the SCENIC+ TF-to-gene network. Red points indicate Bonferroni genome-wide significant genes. Red points indicate nominal (*p* < 0.05) significant genes. Gray points indicate non-significant genes. **(d)** ATAC signal across the *HEYL* locus in young, transition and post-menopausal samples, shading the age-closing −5.5 kb upstream element (green) and ANM-TF motif locations. **(e)** JASPAR2020 motifs for ANM-associated TF families enriched in the element (*SOX*, *PPARG*, *FOXO*), with corresponding motif instances and strands shown. **(f)** Sample-level accessibility of the −5.5 kb element versus donor age (Spearman ρ = −0.79, p = 8 × 10^-4^). **(g)** MAGMA gene-level association (ANM, Ruth 2021) for *HEYL* upstream TFs.

Notably, *HEYL* also showed a gene-level association with ANM (**Fig. 5b**). We next investigated whether predicted upstream regulators of *HEYL* were associated with ANM genetics. Among 81 predicted regulators, five genes—*BRCA1*, *FOXO1*, *PPARG*, *ZNF580* and *ZSCAN31*—showed significant gene-level associations with ANM, with SOX6 exhibited a nominal association (**Fig. 5c**; **Supplementary Fig. 8c**).

We further examined age-associated chromatin accessibility changes at the *HEYL* locus. Accessibility of a candidate regulatory region approximately 5.5 kb upstream of *HEYL* was negatively correlated with donor age (**Fig. 5d.f**). Motif analysis of this region identified putative binding sites for *PPARG*, *SOX* and *FOXO* family transcription factors (**Fig. 5e**). Among these, *PPARG* is an established GC-enriched nuclear receptor involved in steroidogenesis and luteinisation ^42^ The *PPARG* locus also exhibited age-associated chromatin accessibility changes (**Supplementary Fig. 8d**) and gene-level genetic association with ANM(**Fig. 5g; Supplementary Fig. 8e**). Both the canonical and alternative promoters of *PPARG* showed progressively reduced accessibility with age in GC (**Supplementary Fig. 8f,g**).

Together, these findings link reproductive aging-associated genetic signals to predicted regulatory networks and age-sensitive chromatin regions in ovarian cells, highlighting potential regulatory relationships involving *HEYL* and its upstream regulators in GCs.

### Spatial mapping of GC programs and reproductive aging genetics

We next examined the spatial organization of ovarian cell populations and their associated transcriptional programs using spatial transcriptomics (**Fig. 6a**). Although tissue sections were profiled from all three reproductive stages, the transition-stage section contained only five granulosa cell (GC)-assigned bins and was therefore excluded from subsequent GC-focused analyses. We thus focused on sections from young and postmenopausal donors (**Supplementary Table 1**). Cell-type annotations transferred from the scRNA-seq reference were supported by the spatial distributions of canonical markers. GC-assigned bins localized to follicle-associated regions in the young ovary, surrounded by other ovarian cell populations (**Fig. 6a,b**). Compared with the young section, the postmenopausal section showed a lower proportion of GC-assigned bins and altered spatial distributions of ovarian cell populations (**Fig. 6b,c**). Spatial neighborhood analysis further revealed greater GC-macrophage adjacency in the postmenopausal section (**Fig. 6d**).

**Figure 6:**
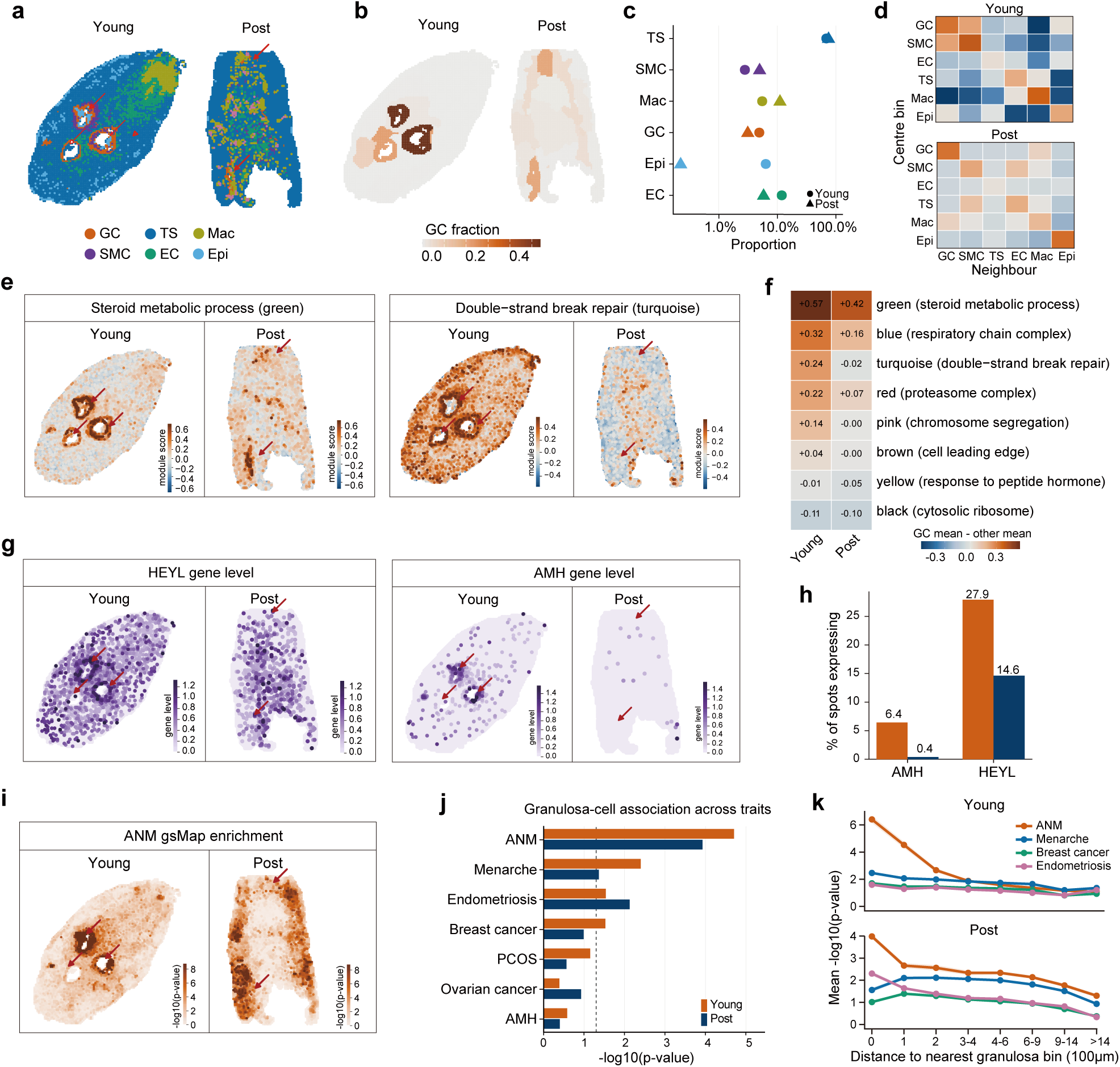
Spatial transcriptomics localizes age-associated changes and ANM genetic signal to granulosa cells in the follicle. **(a)** Spatial cell-type maps of young and post-menopausal ovarian sections. Cell types were assigned by label transfer from the single-cell RNA-seq reference. **(b)** GC fraction per spatial bin in the young and post-menopausal sections. **(c)** Proportion of spatial bins assigned to each cell type in the young and post-menopausal sections. **(d)** Neighborhood co-occurrence of cell types, shown as log2(observed/expected) number of adjacent bin pairs, where the expected numbers are derived from cell-type frequencies. **(e)** Spatial module scores of the GC co-expression modules green (steroid metabolic process) and turquoise (double-strand break repair) defined in Figure 3k. **(f)** Follicle restriction of the eight GC co-expression modules, shown as the mean module score in GC bins minus the mean in all other bins. **(g)** Spatial expression of *HEYL* and *AMH* gene. **(h)** Percentage of bins expressing *AMH* and *HEYL* in each section. **(i)** Per-bin ANM gsMap association (−log10(*p*-value)). ANM GWAS summary data was from Ruth et al. 2021. **(j)** gsMap association of GC bins across ovarian traits, shown as −log10 of the median per-bin *p* value. **(k)** Mean gsMap −log10(*p*-value) of each trait as a function of distance to the nearest GC bin.

We further mapped GC-associated co-expression modules identified by single-cell analysis onto the spatial transcriptomic profiles (**Fig. 3m**; **Fig. 6e,f**). The steroid metabolic module (green) showed higher scores in follicle-associated regions of the young section, whereas its spatial distribution appeared more diffuse in the postmenopausal section. The DNA double-strand break repair module (turquoise) showed lower scores across the postmenopausal section, consistent with its declining activity along the GC pseudotime trajectory (**Fig. 6e**). At the gene level, *AMH* expression was concentrated in follicle-associated regions of the young section but was rarely detected in the postmenopausal section. Similarly, *HEYL* showed a lower proportion of expressing bins in the postmenopausal section compared with the young section (**Fig. 6g,h**). These observations provide spatial context for the GC-associated transcriptional changes identified by single-cell analysis.

We next mapped genetic associations with reproductive traits into the spatial transcriptomic profiles using gsMap ^19^ (Fig. **6i-k**). Among the traits examined, ANM showed the strongest genetic association enrichment in GC-assigned bins in both sections (**Fig. 6j**). In the young section, ANM-associated enrichment was highest in GC-assigned bins and decreased with increasing distance from the nearest GC bin (**Fig. 6k**). Together, these findings highlight the spatial enrichment of ANM-associated genetic signals in GC-rich ovarian regions, complementing the cell-type-specific genetic associations identified by single-cell analysis.

### *HEYL* restrains senescence-associated changes in granulosa cells

To investigate the role of *HEYL* in ovarian aging, we first examined its expression in human ovarian tissues. Immunofluorescence staining revealed predominant localization of *HEYL* in FSHR-positive GC regions (**Fig. 7a; Supplementary Table 8**). *HEYL* mRNA expression was lower in GCs from women aged ≥ 35 years than in those from younger women (**Fig. 7b**). Similarly, *Heyl* expression was reduced in primary mouse GCs isolated from 24-month-old mice compared with those from 3-month-old mice (**Fig. 7c**). In human granulosa-like KGN cells, H_2_O_2_ treatment reduced *HEYL* protein abundance while increasing p21 and p53 protein levels (**Fig. 7d**). These results link reduced *HEYL*/*Heyl* expression with GC aging and oxidative stress-induced senescence-associated changes.

**Figure 7.**
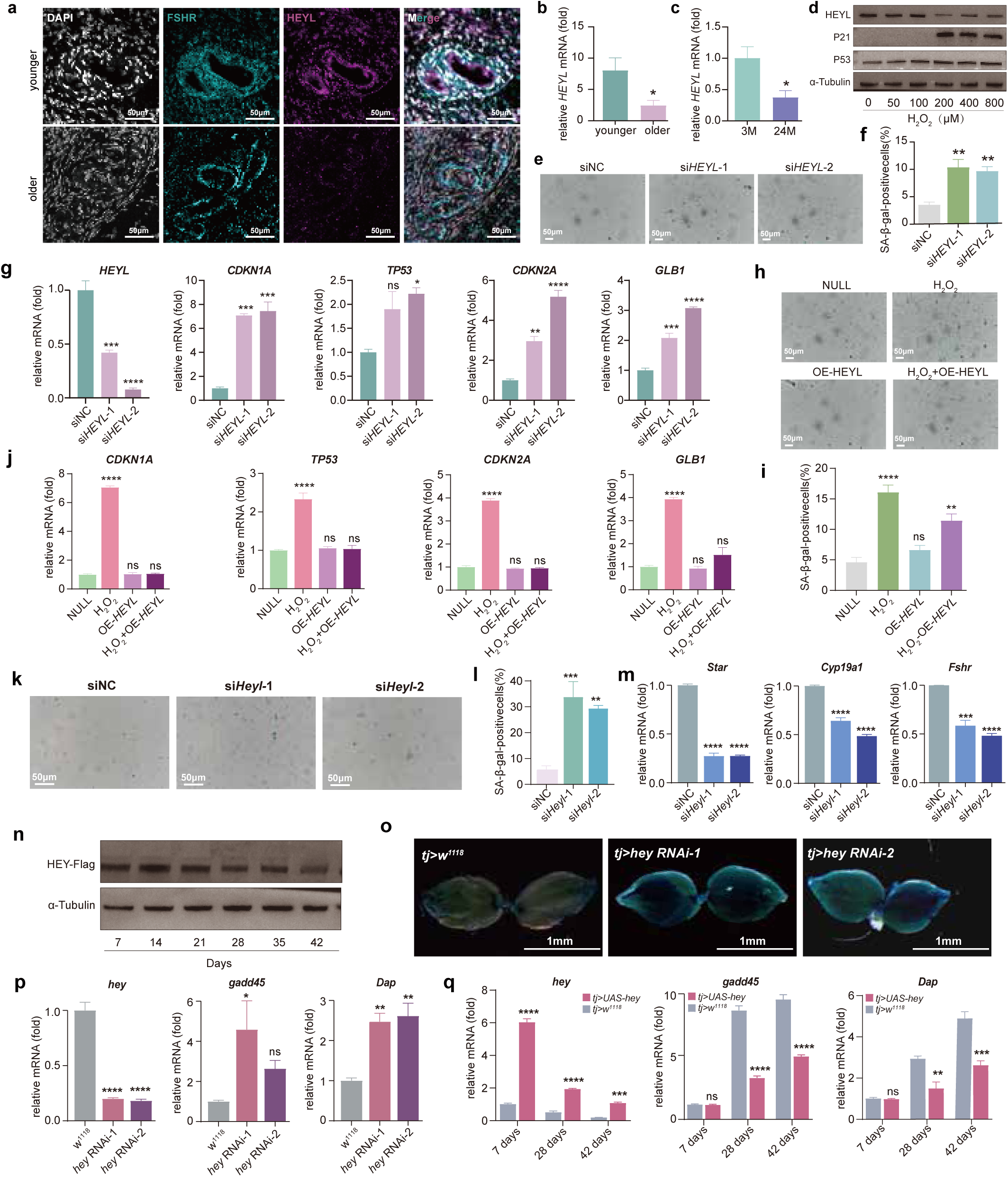
*HEYL* expression declines with ovarian aging and attenuates senescence-associated changes in granulosa cells. **(a)** Representative immunofluorescence images of ovarian sections from younger (<35 years) and older (≥35 years) women stained for FSHR (cyan), HEYL (magenta), and DAPI (white). **(b)** qRT-PCR analysis of relative *HEYL* mRNA levels in granulosa cells from younger and older women. **(c)** Relative *Heyl* mRNA levels in primary granulosa cells isolated from 3- and 24-month-old mice. **(d)** Western blot analysis of HEYL, p21, and p53 in KGN cells treated with increasing concentrations of H_2_O_2_ (0-800 μM). α-Tubulin served as the loading control. **(e, f)** Representative SA-β-gal staining images **(e)** and quantification of SA-β-gal-positive KGN cells **(f)** following *HEYL* knockdown using two independent siRNAs. **(g)** qRT-PCR analysis of *HEYL* and the senescence-associated genes *CDKN1A*, *TP53*, *CDKN2A*, and *GLB1* in KGN cells following *HEYL* knockdown. **(h, i)** Representative SA-β-gal staining images **(h)** and quantification of SA-β-gal-positive KGN cells **(i)** following *HEYL* overexpression with or without H_2_O_2_ treatment. **(j)** qRT-PCR analysis of *CDKN1A*, *TP53*, *CDKN2A*, and *GLB1* under the same treatment conditions. **(k, l)** Representative SA-β-gal staining images **(k)** and quantification of SA-β-gal-positive primary mouse granulosa cells **(l)** following *Heyl* knockdown using two independent siRNAs. **(m)** qRT-PCR analysis of *Star*, *Cyp19a1*, and *Fshr* expression in primary mouse granulosa cells following *Heyl* knockdown. **(n)** Western blot analysis of Flag-tagged Hey protein in *Drosophila* ovaries at the indicated ages (7-42 days). α-Tubulin served as the loading control. **(o)** Representative images of SA-β-gal staining in ovaries from control (*tj>w^1118^*) and ovarian somatic cell-specific *hey* knockdown flies (*tj>hey RNAi-1* and *tj>hey RNAi-2*). **(p)** qRT-PCR analysis of *hey* and the aging- and stress-associated genes *gadd45* and *Dap* in ovaries following ovarian somatic cell-specific *hey* knockdown. **(q)** qRT-PCR analysis of *hey*, *gadd45*, and *Dap* in ovaries from control (*tj>w^1118^*) and *hey*-overexpressing (*tj>UAS-hey*) flies at 7, 28, and 42 days of age. Data are presented as mean ± SEM. *p* < 0.05 (*), *p < 0.01* (**), *p < 0.001* (***) and *p* < 0.0001 (****); ns, not significant. OE, overexpression; SA-β-gal, senescence-associated β-galactosidase; SASP, senescence-f-associated secretory phenotype.

We next examined the effects of *HEYL* depletion and overexpression in KGN cells. Knockdown of *HEYL* with two independent siRNAs increased the proportion of senescence-associated β-galactosidase (SA-β-gal)-positive cells (**Fig. 7e, f**), accompanied by elevated p21 and p53 protein levels (**Supplementary Fig. 9a**). Both siRNAs increased the expression of the senescence-associated genes, including *CDKN1A*, *CDKN2A*, and *GLB1*, whereas *TP53* expression was significantly increased only with si*HEYL*-2 (**Fig. 7g**). *HEYL* knockdown also increased the expression of senescence-associated secretory phenotype (SASP)-related genes, including *IL6*, *CXCL8*, and *SERPINE1* (**Supplementary Fig. 9b**). In contrast, *HEYL* overexpression attenuated H_2_O_2_-induced senescence-associated changes, reducing the proportion of SA-β-gal-positive cells, although levels remained above those of untreated controls (**Fig. 7h, i**). *HEYL* overexpression also reduced p21 and p53 protein levels (**Supplementary Fig. 9c**) and decreased the expression of *CDKN1A*, *TP53*, *CDKN2A*, and *GLB1* to levels not significantly different from those in untreated control (**Fig. 7j**). These results suggest that *HEYL* limits senescence-associated changes in human granulosa-like cells.

We further examined the role of *Heyl* in primary mouse GCs. *Heyl* knockdown increased SA-β-gal positivity (**Fig. 7k,l**) and the expression of the senescence-associated genes, including *Cdkn1a*, *Cdkn2a*, and *Glb1*, whereas *Trp53* expression remained unchanged (**Supplementary Fig. 9e**). SASP-related genes, including *Il6*, *Cxcl1*, and *Serpine1,* were also upregulated (**Supplementary Fig. 9f**). Moreover, *Heyl* knockdown reduced the expression of GC functional genes, including *Star*, *Cyp19a1*, and *Fshr* (**Fig. 7m**). Conversely, *Heyl* overexpression overexpression attenuated H_2_O_2_-induced expression of senescence-associated and SASP-related genes (**Supplementary Fig. 9g,h**), while maintaining *Star*, *Cyp19a1*, and *Fshr* expression at levels comparable to untreated controls(**Supplementary Fig. 9i**).

To assess whether these effects extend across species, we examined *Hey*, the *Drosophila* orthologue of *HEYL*, in the fly ovary. Western blot analysis revealed an age-associated decline in tagged *Hey* protein abundance (**Fig. 7n**). Ovarian somatic cell-specific *hey* knockdown increased ovarian SA-β-gal stainingand the expression of the aging- and stress-associated gene *Dap* (**Fig. 7o,p**). *Gadd45* expression was also increased, although this increase reached statistical significance in only one of the two RNAi lines (**Fig. 7p**). Expression of *puc* and *Socs36E* was also elevated following *hey* depletion (**Supplementary Fig. 9j**). In contrast, ovarian somatic cell-specific *hey* overexpression reduced *gadd45* and *Dap* expression at 28 and 42 days of age (**Fig. 7q**), as well as *puc* expression at 42 days, while *Socs36E* expression was not significantly altered (**Supplementary Fig. 9k**). These results suggest that *Hey* also modulates aging-associated molecular changes in the *Drosophila* ovary. Together, these findings establish *HEYL* as a regulator of senescence-associated changes in GCs, with concordant effects of *Hey* in *Drosophila* suggesting a potentially conserved role in ovarian aging.

## Discussions

Here, we provided a stage-resolved view of human ovarian aging by integrating single-cell transcriptomic, chromatin accessibility and spatial transcriptomic profiles. By distinguishing the menopausal transition from postmenopausal stages, our study extends previous ovarian atlas that largely examined broad age-defined cohorts^11–13^. We identified extensive GC remodelling during reproductive aging, with a reduced proportion of pGCs and an increased proportion of atretic GCs already evident during the transition stage. Notably, transition-stage GCs showed distinct transcriptional changes, including reduced expression of DNA damage-response genes and elevated stress-response programs. At the chromatin level, most age-associated changes occurred outside promoter regions. In GCs, regions losing accessibility were enriched for nuclear receptor motifs associated with steroidogenic and metabolic regulation, whereas regions gaining accessibility were enriched for stress-response AP-1 motifs. These results suggest coordinated remodelling of GC identity-associated and stress-response programs during ovarian aging, with distinct molecular changes emerging during the menopausal transition. Our findings extend previous reports implicating mTOR signaling, *FOXP1* and oxidative phosphorylation in ovarian aging ^11,12^.

Our integrative analyses further identified *HEYL* as a candidate regulator of ovarian aging, supported by age-associated chromatin accessibility changes and GC-enriched regulatory networks. *HEYL* is a canonical downstream target of Notch signaling ^43^, which has established roles in follicular development and GC function ^44,45^. Given that oocyte-derived Notch ligands contribute to Notch signaling activation in GCs during follicular development, the age-associated decline in *HEYL* expression may reflect altered oocyte-GC communication accompanying follicular depletion. Our functional findings further implicate *HEYL* in maintaining GC function and limiting senescence-associated changes, with concordant effects in *Drosophila* suggesting a potentially conserved role in ovarian aging. The induction of inflammatory genes following *HEYL* depletion also raises the possibility that its regulatory effects extend to the follicular microenvironment, although this remains to be investigated.

Integration with reproductive aging genetics provided additional support for the potential involvement of *HEYL*-associated regulatory networks in ovarian aging. Several upstream regulators of *HEYL*, including *PPARG* and *FOXO1*, showed gene-level genetic associations with ANM. These associations are consistent with the established roles of *PPARG* in GC steroidogenesis and luteinization ^42,46^ and FOXO family members in follicle survival and atresia ^47^. Furthermore, age-associated reductions in chromatin accessibility at the *HEYL* and *PPARG* loci, together with putative PPARG- and FOXO-family binding motifs near *HEYL*, suggest potential regulatory relationships linking these factors to GC aging-associated programs. Although these interactions remain to be functionally established, our findings raise the possibility that, beyond the oocyte DNA damage-response pathways emphasized by previous GWAS ^6,48^, genetic variation associated with menopause timing may also involve regulatory programs in GCs.

Several limitations should be considered. First, ovarian tissues were obtained from surgical donors with varying clinical indications and sampling sites: mainly cortical biopsies for young donors and mainly oophorectomy specimens for older donors. These differences may have contributed to the observed changes in cell-type composition, although major cell populations and their markers were consistent across donors and modalities. Second, tissue dissociation and filtration may preferentially exclude large or fragile cells, including oocytes, which were not captured in our atlas. Third, scRNA-seq and scATAC-seq were performed on separate cells, and the inferred regulatory relationships require further validation through paired multi-omic profiling or direct binding and perturbation assays. Fourth, the limited abundance of GCs in postmenopausal ovaries and the scarcity of GC-assigned bins in transition-stage spatial samples constrained stage-specific analyses. Spatial comparisons were also based on one tissue section per reproductive stage, limiting assessment of inter-individual variability. Finally, although functional experiments in human granulosa-like cells, primary mouse GCs and *Drosophila* provided support for the regulatory role of *HEYL*, its effects on ovarian function and reproductive lifespan in mammals in vivo remain to be established.

In summary, our study provides a stage-resolved multi-omic atlas of human ovarian aging, revealing extensive GC remodelling across reproductive stages and identifying *HEYL* as a regulator of senescence-associated changes. These findings advance our understanding of the cellular and regulatory changes accompanying the menopausal transition and provide a foundation for investigating mechanisms underlying ovarian functional decline.

## Methods

### Human ovarian tissue collection and ethics

Human ovarian tissues were obtained from participants undergoing ovarian biopsy or oophorectomy at Women’s Hospital, Zhejiang University School of Medicine. The study was registered at ClinicalTrials.gov (NCT06067529) and approved by the Ethics Committee of Women’s Hospital, Zhejiang University School of Medicine (IRB-20220393-R). Written informed consent was obtained from all participants before sample collection.

Ovarian tissue donors were collected and assigned to young, transition or post groups according to age, menstrual history and serum follicle-stimulating hormone (FSH) levels. The young group comprised reproductive-age women with regular menstrual cycles, basal FSH <10 IU/L and preserved ovarian reserve. The transition group comprised non-menopausal women aged 40-55 years with evidence of reproductive aging. The post group comprised women with amenorrhoea for more than 12 months and FSH >25 IU/L. Detailed donor-level clinical information, including age, diagnosis, hormone measurements and sequencing modality, is provided in **Supplementary Table 1**.

Ovarian cortical biopsies were obtained from young donors when clinically feasible, whereas tissues from transition and post donors were obtained mainly from oophorectomy specimens. Surgical indications included benign ovarian or gynaecological diseases and selected early-stage non-ovarian malignancies or tumours of uncertain malignant potential. Only macroscopically uninvolved ovarian tissue was collected, and final pathological examination confirmed the absence of ovarian tumour involvement.

scRNA-seq and scATAC-seq were each performed on five donors per group. Most donors were profiled by both modalities from the same specimen, whereas a small number of samples were profiled by a single modality because of limited tissue availability or library-quality failure. Spatial transcriptomics was performed on one donor per group. Fresh ovarian tissue was placed in pre-cooled preservation medium within 15 min after surgical removal and transported under cold-chain conditions for downstream processing.

### Human granulosa cell collection and ethical approval

Human granulosa cells were obtained from women undergoing in vitro fertilization and embryo transfer (IVF-ET) for male-factor infertility at the Women’s Hospital, School of Medicine, Zhejiang University. The study was approved by the Ethics Committee of the Women’s Hospital, Zhejiang University School of Medicine (approval No. IRB-20230043-R), and written informed consent was obtained from all participants before sample collection. A total of 29 women were enrolled and divided into two age groups: < 35 years (n = 14) and ≥ 35 years (n =15). Follicular fluid was collected during transvaginal ultrasound-guided oocyte retrieval following controlled ovarian stimulation. Granulosa cells were isolated from the collected follicular fluid and subsequently used for RNA extraction and quantitative real-time PCR (qRT-PCR) analysis. Detailed clinical characteristics and exclusion criteria for the participants are provided in Supplementary Table 2.

### Single-cell RNA sequencing

scRNA-seq libraries were prepared by Beijing SeekGene BioSciences Co., Ltd (Beijing, China) using the SeekOne Digital Droplet Single Cell 3′ Library Preparation Kit (SeekGene, K00202) according to the manufacturer’s instructions. Libraries were sequenced on a DNBSEQ-T7 platform.

Sequencing reads were aligned to the human reference genome GRCh38 and quantified with SeekSoulTools (SeekGene) to generate gene-cell UMI count matrices. Downstream analyses were performed in Seurat (v4.4.0) ^49^. Low-quality cells were removed (< 500 UMIs, < 200 detected genes, > 10% mitochondrial content). After quality control, 183,394 cells and 31,081 genes were retained. Counts were log-normalized, and highly variable genes were selected, scaled and reduced by principal component analysis. Donor effects were corrected with Harmony (v1.2.4) ^50^. The Harmony embedding was used for UMAP and shared-nearest-neighbour clustering. Clusters were annotated to ovarian cell types using canonical markers.

### Single-cell ATAC sequencing

scATAC-seq libraries were generated by Hangzhou Astrocyte Technology Co., Ltd (Hangzhou, China) using the Chromium Next GEM Single Cell ATAC Reagent Kits (10x Genomics; PN-1000110, PN-1000156, PN-1000084) according to the manufacturer’s instructions. scATAC-seq libraries were sequenced on an Illumina NovaSeq X Plus platform.

Fragment files from Cell Ranger ATAC (v2.1.0; GRCh38) were analysed with Signac (v1.17.0) ^51^ and Seurat (v4.4.0) ^49^. A consensus peak set was generated by merging the per-library Cell Ranger peak calls (GenomicRanges::reduce) and retaining peaks 20-10,000 bp in length. Peaks were quantified in each library with FeatureMatrix for barcodes with >500 passed-filter fragments. Peaks on non-standard chromosomes were removed, and genes were annotated with EnsDb.Hsapiens.v86. Per-nucleus quality metrics were computed with Signac ^51^: nucleosome signal, TSS enrichment score, fraction of fragments in peaks (FRiP) and fraction of fragments in ENCODE hg38 blacklist regions. Nuclei were retained if they had 3,000-30,000 fragments in peaks, TSS enrichment > 3, nucleosome signal < 4, FRiP > 15% and blacklist fraction < 0.05. This yielded 129,936 nuclei and 330,160 peaks. Data were normalized by term frequency-inverse document frequency (TF-IDF) using all peaks and reduced by singular value decomposition (latent semantic indexing, LSI). Donor effects were corrected with Harmony on the LSI embedding. Harmony components 2-30 were used for UMAP and shared-nearest-neighbour clustering. Gene-activity scores were computed with GeneActivity and log-normalized, with the median gene-activity count per nucleus as the scale factor. Cell types were annotated at the cluster level based on gene-activity scores of canonical marker genes for each ovarian cell type.

### Concordance between scRNA-seq and scATAC-seq annotations

Marker genes used for scRNA-seq annotation were evaluated in the corresponding scATAC-seq cell types. For each marker gene, chromatin accessibility was quantified within its promoter (±500 bp around the annotated TSS) and aggregated across nuclei of each scATAC-seq cell type. Values were scaled across cell types and shown as a heatmap (**Fig. 1f**). Overlap between the two modalities was quantified with the Jaccard index between cell-type-specific gene sets. The scRNA-seq sets were differentially expressed genes. The scATAC-seq sets were differentially active genes, identified from gene-activity scores (**Fig. 1g**).

### Cell-type composition analysis

Cell-type proportion changes with age were assessed using *sccomp* (v2.1.23) ^20^. Cell-type abundances per donor were modeled using *sccomp_estimate*, followed by significance testing with *sccomp_test*. Two models were fitted, one with continuous age (∼ age) and one with age group (∼ age group; young as reference). In both, the same term was used in the composition formula and in the variability formula. Significance was defined by posterior credible intervals excluding zero.

### Differential expression analysis

Age-associated gene expression changes were evaluated separately within each cell type using NEBULA (v1.5.6) ^22^. Raw UMI counts were modeled with a negative-binomial mixed-effects model (NBGMM), with donor as a random effect and total UMI count per cell as an offset. Age was modeled either as continuous age in years or as age group. *P* values were adjusted using the Benjamini-Hochberg (BH) procedure within each cell type. Differentially expressed genes (DEGs) were defined as FDR < 0.05 and |log fold-change| > log(1.1). Gene Ontology (GO) biological-process enrichment of up- and down-regulated DEGs was tested with a hypergeometric test against MSigDB C5 GO:BP gene sets.

### Aging-gene scoring and trajectory clustering

A curated aging-associated gene set from GenAge ^23^ was scored in individual cells using Seurat AddModuleScore. Differences in aging scores across cell types were assessed using the Wilcoxon rank-sum test. Within GCs, the expression of GenAge genes was averaged per age group (young, transition and post), and the genes were grouped using unsupervised clustering. Five expression-pattern clusters were identified and each was tested for GO enrichment. GO enrichment analyses of module genes were performed using clusterProfiler (v4.18.4)^52^.

### Co-expression network analysis

Cell-type-specific co-expression networks were constructed using hdWGCNA (v0.4.8) ^24^. For each cell type, the top 2,000 highly variable genes were selected and aggregated into donor-aware metacells (k = 25, max_shared = 10). Signed weighted gene co-expression networks were generated using the minimum soft-thresholding power achieving a scale-free topology fit of *r*^2^ > 0.85. Modules were identified using ConstructNetwork (deepSplit = 4, minModuleSize = 50, mergeCutHeight = 0.2), and module eigengenes were calculated while controlling for donor effects. Modules were summarized using the top 25 hub genes and scored across cells using both AddModuleScore and UCell. Associations between module activity and age were evaluated using Spearman correlation with BH correction. A granulosa-cell-specific network was constructed using the same framework, except with mergeCutHeight = 0.5. GO and pathway enrichment analyses of module genes were performed using clusterProfiler (v4.18.4) ^52^.

### Granulosa-cell subtype identification and trajectory analysis

GC were re-clustered using Seurat (resolution = 0.1) and annotated based on established marker genes as pGC, cumulus, mural, and atretic GC populations. Differentiation trajectories were reconstructed using Monocle3 (v1.4.26, https://cole-trapnell-lab.github.io/monocle3/). A UMAP embedding was computed from the first five Harmony-corrected components. The data were converted to a cell_data_set object, cells were clustered with cluster_cells, and a principal graph was learned across all cells with learn_graph (use_partition = FALSE). Cells were ordered along the trajectory with order_cells. The root was the principal-graph node closest to the largest number of cells from young donors. For downstream comparisons, pseudotime was divided into four intervals. Pseudotime distributions were compared between age groups using the Wilcoxon rank-sum test. Within each pseudotime interval, GC subtype composition was compared between age groups using Fisher’s exact test. The differentiation potential of each cell was estimated using CytoTRACE ^28^.

### Cell-cell communication analysis

Intercellular communication was inferred separately for the young, transition and post groups using CellChat (v2.2.0) ^53^ with log-normalized expression data. Ligand-receptor interactions were drawn from the full human CellChatDB database. Overexpressed ligands, receptors and interactions were identified in each cell type. Communication probabilities were then computed with the trimean method, without projection onto a protein-protein interaction network. Interactions involving cell groups with fewer than 10 cells were removed. Communication probabilities were aggregated to the signaling-pathway level and summarized as networks of interaction number and interaction strength. Network centrality analysis identified the dominant senders and receivers within each group.

### Differential chromatin accessibility and motif analysis

DARs were identified within each cell type using Seurat (v.4.4.0)^49^ logistic regression (FindMarkers, test.use = “LR”, with the number of fragments in peaks as a latent variable). Peaks accessible in at least 5% of nuclei in either group and with |log2FC| ≥ 0.25 were tested. Analyses were restricted to cell types with more than 50 nuclei and more than 10 nuclei per group. DARs were defined by Bonferroni-adjusted p < 0.05. DAR peaks were annotated to genomic features and nearest genes using ChIPseeker (v1.46.1) ^54^ with TxDb.Hsapiens.UCSC.hg38.knownGene. Promoters were defined as 3 kb around the TSS.

For motif analyses, donor-level pseudobulk profiles were generated for each cell type by summing fragment counts across nuclei. Only donors contributing ≥ 50 nuclei were included ( ≥ 10 for GCs and rare cell types), and each cell type required at least 3 donors per age group. Peaks were filtered with edgeR filterByExpr (min.count = 5, min.total.count = 15) and TMM-normalized. Motif occurrences in peaks were identified using motifmatchr and the JASPAR2020 CORE human collection. Motif enrichment among gained or lost DARs was assessed using one-sided Fisher’s exact tests, relative to all peaks tested in the same cell type.

Motif activity was quantified using chromVAR (v1.32.0) ^34^. Deviation scores were computed for non-overlapping peaks after GC-bias correction (BSgenome.Hsapiens.UCSC.hg38). Scores were then averaged per donor within each cell type (cell types with more than 500 nuclei; donors with more than 50 nuclei). Associations with continuous age and with age group were tested using limma ^55^.

### *HEYL* regulatory network analysis

Enhancer-driven regulons (eRegulons) were inferred using SCENIC+ ^35^. eRegulon activity was quantified as AUC scores in metacells. Cell-type specificity was defined as the mean z-scored AUC per cell type.

Co-accessibility between the *HEYL* distal element (peak 11) and surrounding peaks (chr1:39,603,000-39,660,000) was estimated as the Pearson correlation of binarized accessibility across all nuclei. Empirical *p* values were derived from 2,000 randomly sampled peaks outside the locus.

*HEYL* upstream and downstream genes were defined from the GRNBoost network. These targets were tested for enrichment in the GO biological process and KEGG pathway enrichment. The background was all genes in the GRNBoost target space (29,738 genes), and significance was BH-adjusted *p* < 0.05.

### Gene enrichment analysis of reproductive-trait GWAS summary data

Gene-level association statistics were computed with MAGMA v1.10 using the SNP-wise mean model. SNPs were mapped to genes within a ±10 kb window (NCBI37.3 gene locations), and the 1000 Genomes Phase 3 European panel was used as the LD reference. Trait gene sets include MAGMA FDR < 0.05 genes for large-scale traits (ANM Ruth 2021/Day 2018, menarche, breast cancer) and the top 1,000 genes ranked by ZSTAT for smaller sets (endometriosis, ovarian cancer, AMH, PCOS).

Enrichment of trait genes among age-associated DEG and DAR-linked gene sets was assessed using one-sided Fisher’s exact tests. The background for each cell type was the set of genes tested by NEBULA in that cell type, and analyses required more than 20 genes per set and more than 5 overlapping genes.

### Spatial transcriptomic profiling and preprocessing

Ovarian sections were profiled by Stereo-seq (Annoroad) and processed with the SAW pipeline. Counts were aggregated into Bin200 units (100 × 100 µm), and each bin, rather than each cell, was treated as the unit of analysis. Bins were annotated by label transfer from the scRNA-seq reference. Anchors were identified with FindTransferAnchors (LogNormalize, PCA, dims 1-30, k.anchor = 30), and each bin was assigned the label with the highest TransferData score (k.weight = 20). The perimenopausal section contained only five GC-assigned bins and was excluded from GC-focused analyses.

To define spatially contiguous domains, principal components (30 PCs from 2,000 highly variable genes) were smoothed over each bin and its four adjacent bins. Bins were then clustered with Leiden clustering (15-nearest-neighbour graph, resolution 0.5). Cell-type co-occurrence was quantified across adjacent bin pairs (rook adjacency, four neighbours) as log2((observed + 1)/(expected + 1)). Expected counts were derived from the marginal cell-type frequencies.

### Module scoring and marker genes

GC co-expression modules from the hdWGCNA^24^ analysis were scored in each bin using the top 10 hub genes per module, with an AddModuleScore approach. Module enrichment in GC-assigned bins was expressed as the difference in mean score between GC and non-GC bins. For *AMH* and *HEYL*, we report the percentage of bins with detectable expression (non-zero counts) and the log_2_ ratio of mean counts per 10,000 (CP10K) between GC and non-GC bins.

### Spatial mapping of GWAS associations

Trait associations were mapped onto each section with gsMap (v1.73.8) ^19^ using the GWAS summary statistics for reproductive traits described above. Per-bin association *p* values were summarized for each cell type as the median *p* value. For distance profiles, the distance from each bin to the nearest GC-assigned bin was computed. Mean per-bin −log_10_(*p*-value) was then plotted in distance intervals.

### Mouse husbandry

Female C57 mice were maintained under specific-pathogen-free conditions at 22 ± 2°C under a 12-h light/12-h dark cycle, with ad libitum access to standard chow and water.

#### *Drosophila* stocks and husbandry

The following *Drosophila melanogaster* stocks were used: PBac{Hey-GFP.FPTB}VK00033 (BDSC no. 64826), the *Hey* RNAi line THU2274 (Tsinghua Fly Center), M{UAS-Hey.ORF.3xHA.GW}ZH-86Fb (FlyORF no. F000566), and the ovarian somatic cell driver *tj-GAL4/CyO*, which was maintained in our laboratory.

Flies were maintained on standard fly food under conventional culture conditions at 25℃ with a 12 h light/ 12h dark cycle. Newly enclosed female flies were collected within 6 h of eclosion and housed with males. Flies were maintained per vial and transferred to fresh food every 2 days.

### *Drosophila* aging and ovary collection

Female flies were collected within 6 h after eclosion, co-housed with males, and maintained at a density of 10 females and 5 males per vial. Flies were transferred to fresh food every 2 days until the indicated ages. At 7, 14, 21, 28, 35, or 42 days after eclosion, ovaries were dissected in Schneider’s *Drosophila* Medium (Gibco, Thermo Fisher Scientific, Cat. No. 21720024).

For age-dependent analysis of Hey protein abundance, ovaries were collected from flies carrying PBac{Hey-GFP.FPTB}VK00033. Tagged Hey protein was detected using an anti-FLAG antibody.

#### Collection of ovaries from young and aged mice

Young and aged female C57 mice were euthanized at 12 weeks and 24 months of age, respectively. Estrous cyclicity was monitored by daily vaginal cytology for 7 consecutive days before tissue collection. Young mice were euthanized during diestrus to minimize variation associated with estrous-cycle-dependent hormonal fluctuations, whereas aged mice were confirmed to be acyclic based on vaginal cytology. All animals were euthanized during the same time window of the light phase. Ovaries were immediately excised, and the surrounding adipose tissue and oviducts were carefully removed under a stereomicroscope. Both ovaries from each mouse were pooled, snap-frozen in liquid nitrogen, and stored at-80°C until RNA extraction.

#### H_2_O_2_-induced oxidative senescence in KGN cells

To optimize the oxidative senescence model, KGN cells were initially exposed to 0, 50, 100, 200, 400, or 800 μM hydrogen peroxide (H_₂_O_₂_). Immediately before use, a fresh 200 mM intermediate H_₂_O_₂_ stock solution was prepared from 30% H_₂_O_₂_. This 1,000× stock solution was subsequently diluted 1:1,000 in complete culture medium to obtain a final H_₂_O_₂_ concentration of 200 μM.

For subsequent experiments, KGN cells were treated with 200 μM H_₂_O_₂_ in a serum-containing medium for 20 min. The H_₂_O_₂_-containing medium was then removed and replaced with a fresh complete medium. Control cells underwent the same medium-replacement procedure without H_₂_O_₂_ exposure.

Cells were collected 48 h after H_₂_O_₂_ treatment for RNA analysis and 60-72 h after treatment for protein analysis. The 200 μM condition was selected because it induced a detectable senescence-associated phenotype without causing extensive cell death.

#### siRNA-mediated *HEYL* knockdown in KGN cells

KGN cells were seeded at a density of approximately 1×10^5 cells/mL and transfected with *HEYL*-targeting siRNAs or a non-targeting control siRNA using Lipofectamine RNAiMAX in Opti-MEM. The final siRNA concentration was 20 nM. Cells were collected 48 h after transfection for RNA extraction and quantitative real-time PCR analysis and 72 h after transfection for protein analysis. *HEYL* knockdown efficiency was evaluated at both the mRNA and protein levels.

The *HEYL-*targeting siRNAs were synthesized by Tsingke Biotechnology and had the following sequences (5′-3′): si*HEYL-1* sense: GAGACCGCAUCAACAGUAG; si*HEYL-*1 antisense: CUACUGUUGAUGCGGUCUC; si*HEYL*-2 sense: GAAUGUGCUGCCCAGUCGA; and si*HEYL*-2 antisense: UCGACUGGGCAGCACAUUC.

#### *HEYL* overexpression in KGN cells

KGN cells were transiently transfected with a pcDNA3.1-based *HEYL* expression plasmid or the corresponding empty pcDNA3.1 vector using Lipofectamine 3000. Equal amounts of plasmid DNA were used in all groups.

The transfection medium was replaced with fresh complete medium 4 h after transfection. At 24 h after transfection, cells were treated with 200 μM H_₂_O_₂_ for 20 min. The H_₂_O_₂_-containing medium was then removed and replaced with fresh complete medium. Cells were collected 48 h after H_₂_O_₂_ treatment for SA-β-galactosidase staining and RNA analysis and 72 h after treatment for protein analysis. *HEYL* overexpression efficiency was verified by quantitative real-time PCR and Western blotting.

#### Isolation and culture of primary mouse granulosa cells

Primary granulosa cells were isolated from 8-week-old female C57 mice. To stimulate follicular development, mice were intraperitoneally injected with 20 IU pregnant mare serum gonadotropin (PMSG; Solarbio, Beijing, China; Cat. No. P9970). Ovaries were collected 48 h after PMSG administration.

After removal of the surrounding adipose tissue and oviducts, granulosa cells were released from antral follicles by puncturing the follicles with a needle. The resulting cell suspension was treated with ACK red blood cell lysis buffer for 60 s, after which the lysis reaction was immediately terminated by adding culture medium. The cell suspension was then passed through a 70 μm cell strainer and centrifuged at 500 × g for 5 min. The cell pellet was resuspended in DMEM/F-12 medium supplemented with 10% fetal bovine serum and 1% penicillin-streptomycin and seeded at a density of 1×10^5^cells/mL.

#### *Heyl* knockdown in primary mouse granulosa cells

Primary mouse granulosa cells were transfected with two independent *Heyl*-targeting siRNAs or a non-targeting control siRNA using Lipofectamine RNAiMAX. The final siRNA concentration was 20 nM.

Cells were collected 48 h after transfection for quantitative real-time PCR analysis and senescence-associated β-galactosidase staining. *Heyl* knockdown efficiency was assessed by quantitative real-time PCR and, when sufficient material was available, further confirmed by Western blotting. Each independent primary granulosa-cell isolation was considered one biological replicate.

The mouse *Heyl*-targeting siRNA sequences were as follows (5′-3′): si*Heyl*-1 sense: GGUUCUAUCUCCAGUCCAU; si*Heyl*-1 antisense: AUGGACUGGAGAUAGAACC; si*Heyl*-2 sense: GCAGAGGGAUCAUAGAGAA; and si*Heyl*-2 antisense: UUCUCUAUGAUCCCUCUGC.

#### *Heyl* overexpression in primary mouse granulosa cells

Primary mouse granulosa cells were transiently transfected with a pcDNA3.1-based mouse *Heyl* expression plasmid or the corresponding empty pcDNA3.1 vector using Lipofectamine 3000 and P3000 Reagent.

The transfection medium was replaced with fresh complete medium 4 h after transfection. At 24 h after transfection, cells were treated with 200 μM H_₂_O_₂_ for 20 min. The H_₂_O_₂_-containing medium was then removed and replaced with a fresh complete medium. Cells were collected 24 h after H_₂_O_₂_ treatment for quantitative real-time PCR analysis of *Heyl*, senescence-associated genes, and granulosa-cell functional genes. Senescence-associated β-galactosidase staining was performed 48 h after H_₂_O_₂_ treatment. *Heyl* overexpression efficiency was verified by quantitative real-time PCR and Western blotting.

### *hey* knockdown in *Drosophila* ovarian somatic cells

To knock down *hey* in ovarian somatic cells, *tj-GAL4/CyO* females were crossed with males carrying *hey* RNAi line THU2274. Female progeny carrying both *tj-GAL4* and the *UAS-Hey* RNAi construct, but lacking the CyO balancer chromosome, were used as the experimental group. Driver-matched controls were generated by crossing the same *tj-GAL4/CyO* line with *w^11^*^18^ flies. Female progeny were collected within 6 h after eclosion and co-housed with males at a density of 10 females and 5 males per vial. At 3 days after eclosion, ovaries were dissected and collected in Schneider’s *Drosophila* Medium.

### *hey* overexpression in *Drosophila* ovarian somatic cells

To overexpress *hey* in *Drosophila* ovarian somatic cells, the *tj-GAL4/CyO* driver line was crossed with the *UAS-hey* (FlyORF, F000566) were used as the experimental group. Driver-matched controls were generated by crossing the same *tj-GAL4/CyO* line with *w^1118^*flies. Ovaries were collected at 7, 28, and 42 days after eclosion for phenotypic analysis.

## Acknowledgements

We thank the tissue donors for making this study possible. We thank Beijing SeekGene BioSciences Co., Ltd (Beijing, China) for assistance in scRNA-seq related experiments. We are grateful to Astrocyte Technology Co., Ltd (Hangzhou, China) for assisting in scATAC-seq related experiments.

This work was supported by National Natural Science Foundation of China (No. 82571881), Zhejiang Provincial Natural Science Foundation of China (LZ25H040001), 4+X Clinical Research Project of Women’s Hospital, School of Medicine, Zhejiang University [ZDFY2022-4XA101 and ZDFY2023-4X201].

## Supplementary Figure Legends

**Supplementary Figure 1: Sample FSH level correlated with age.** Correlation between serum follicle-stimulating hormone (FSH) levels and chronological age (R = 0.58, *p* = 0.04).

**Supplementary Figure 2: Quality control, staging, and cellular composition across age groups. (a,b)** Per-sample quality control metrics for scRNA-seq and scATAC-seq, ordered from young, transition to post-menopausal samples. **(c,d)** Cell-type composition across age groups for scRNA-seq **(c)** and scATAC-seq **(d)**.

**Supplementary Figure 3: GC aging programs and their temporal dynamics. (a,b)** GO enrichment of genes up-regulated **(a)** and down-regulated **(b)** with age in GC (ranked by *p*-value; coloured by functional category). **(c)** Average expression of each of the five aging-gene clusters across young, transition and post-menopausal groups.

**Supplementary Fig. 4: Complete cell-type module set, module functions and age-associated cell-cell communication. (a)** Module specificity across cell types for the complete set of per-cell-type hdWGCNA modules.

**(b)** Cross-cell-type module overlap: Jaccard similarity between module gene sets. **(c)** Aging-module correlation: Spearman correlation between top-25 hub-gene module score (AddModuleScore) and age per cell type × module. ** *p*_adj_ < 0.05, * *p*_adj_ < 0.1, (.) raw *p* < 0.05, + raw *p* < 0.2; n = 15 samples. **(d)** GO enrichment of the smooth-muscle module SMC|M1 and the endothelial module EC|M3. **(e)** CellChat relative information flow: signaling pathways significantly up-(red) or down-regulated (blue) in post-versus young ovaries. **(f)** Differential signaling chord diagrams for COLLAGEN, LAMININ, MHC-I and APP (post *v.s.* young). **(g)** Cell-type contribution (% of nuclei) to each significantly enriched signaling pathway in the post-menopausal group. **(h)** Total number of inferred interactions and overall interaction strength in young, transition and post-menopausal ovaries.

**Supplementary Fig. 5: GC co-expression module function, aging dynamics and hub-gene networks. (a)** Gene Ontology enrichment for each GC hdWGCNA module. **(b)** Module eigengene activity by age group (young/transition/post) for each module. **(c)** Hub-gene co-expression networks for representative modules.

**Supplementary Fig. 6: Genome-wide DAR characterization and transcription-factor-activity dynamics during ovarian aging. (a)** Number of differentially accessible regions (DARs) per cell type for each pairwise age comparison (transition vs young, post vs transition, post vs young; logistic-regression test, Bonferroni-adjusted *p* < 0.05, |log2FC| > 0.1). Bars above/below zero denote gained (up, red) and lost (down, blue) accessibility. **(b)** Transcription-factor motifs enriched in GC age-associated DARs (one-sided Fisher test against the granulosa tested-peak universe; JASPAR2020 CORE motifs). Left, motifs enriched among peaks that lose accessibility with age: nuclear-receptor motifs (ESRRB, ESRRA, NR5A1, NR2F2, NR4A2); right, motifs enriched among peaks that gain accessibility: AP-1/bZIP motifs (FOS, JUN, FOSL and JUNB dimers). **(c)** Cell-type-resolved motif enrichment in age-associated DARs, computed with the same uniform one-sided Fisher test against each cell type’s own tested-peak background (motifmatchr; JASPAR2020 CORE). Top, age-closing peaks; bottom, age-opening peaks; rows grouped by TF family. asterisks = adjusted *p* (* < 0.05, ** < 0.01, *** < 0.001). **(d)** Heatmap of motif-activity change with continuous age (chromVAR deviation regressed on age) for the top union of differentially active transcription factors (n = 40) across cell types. Color denotes the age coefficient of TF motif activity. **(e)** Granulosa-cell TF motif activity (chromVAR log2(fold-change)) versus TF mRNA log2(fold-change) (NEBULA) for post vs young (top) and post vs transition (bottom). Points are colored by fold-change directions. **(f)** Directional concordance between TF motif activity and target-gene expression. For each TF, the log2 ratio of its differentially expressed (DEG) targets changing in the same versus the opposite direction to its motif-activity change is plotted against the motif-activity log2(fold-change) with continuous age.

**Supplementary Fig. 7: *HEYL*-locus accessibility, transcription-factor activity and target pathways. (a)** *HEY*-family transcription-factor motif activity (chromVAR deviation) in GC across young, transition and post-menopausal groups. Violin plots show per-cell motif z-scores with pairwise Wilcoxon rank-sum *p* values. **(b)** Heatmap of the age-associated accessibility change (NEBULA log2(fold-change), young vs post) for all 17 *HEYL*-locus peaks (rows, by genomic position) across cell types (columns). \**p* < 0.05, \*\**p* < 0.01, \*\*\**p* < 0.001. **(c)** KEGG pathway enrichment of *HEYL* SCENIC+ GRNBoost target genes (aging- and reproduction-related pathways). Dot color denotes pathway category and dot size the number of contributing genes.

Supplementary Fig. 8: Ovarian trait gene enrichment in age-remodeled chromatin, the *HEYL* regulatory circuit and the *PPARG* locus. (a) Enrichment of ovarian trait (Breast cancer) genes in age-associated DAR/DEG gene sets across cell types, age contrasts and directions. Extension of Fig. 5a. **(b)** The same enrichment using curated GWAS Catalog trait genes as an independent gene-set source (ANM, AMH, PCOS, menarche, ovarian cancer and breast cancer). **(c)** Venn diagram of *HEYL* upstream transcription factors (GRNBoost, n = 81) and ANM genome-wide-significant genes (MAGMA, Ruth2021 n = 837; Bonferroni *p* < 0.05). **(d)** Heatmap of ATAC log2(fold-change) (strongest age contrast) for the top 30 ANM-associated age-DAR genes across ovarian cell types. **(e)** UpSet plot of shared genes among the ovarian trait (menarche, AMH, PCOS, ANM) gene sets; intersection sizes (top) and per-trait set sizes (right). **(f)** ATAC coverage across the *PPARG* locus (chr3) in young, transition and post-menopausal ovaries, with the 85 ANM genome-wide-significant SNPs and the 2 aging-associated peaks and their annotations indicated. **(g)** Sample-level GC accessibility of the 2 aging-associated peaks, *PPARG* promoter peak (chr3:12,286,468-12,289,549; Spearman *ρ* = −0.70, *p* = 0.005) and alternative promoter peak (chr3:12,304,597-12,305,648; *ρ* = −0.56, *p* = 0.039) versus donor age.

**Supplementary Figure 9: HEYL and Hey modulate senescence-associated gene expression in granulosa cells and Drosophila ovaries. (a)** Western blot analysis of HEYL, p21, and p53 in KGN cells transfected with control siRNA (siNC) or two independent *HEYL*-targeting siRNAs (siHEYL-1 and siHEYL-2). α-Tubulin served as the loading control. **(b)** qRT-PCR analysis of the SASP-related genes *IL6*, *CXCL8*, and *SERPINE1* in KGN cells following *HEYL* knockdown. **(c)** Western blot analysis of p21 and p53 in KGN cells with or without *HEYL* overexpression and H_₂_O_₂_ treatment. α-Tubulin served as the loading control.

**(d)** qRT-PCR analysis of *IL6*, *CXCL8*, and *SERPINE1* in KGN cells under the same treatment conditions. **(e, f)** qRT-PCR analysis of *Heyl* and the senescence-associated genes *Cdkn1a*, *Trp53*, *Cdkn2a*, and *Glb1* (e), and the SASP-related genes *Il6*, *Cxcl1*, and *Serpine1* (f) in primary mouse granulosa cells following *Heyl* knockdown using two independent siRNAs. **(g-i)** qRT-PCR analysis of *Heyl*, *Cdkn1a*, *Trp53*, *Cdkn2a*, and *Glb1* (g), the SASP-related genes *Il6*, *Cxcl1*, and *Serpine1* (h), and the granulosa cell functional genes *Star*, *Cyp19a1*, and *Fshr* (i) in primary mouse granulosa cells with or without *Heyl* overexpression and H_₂_O_₂_ treatment. **(j)** qRT-PCR analysis of the aging- and stress-associated genes *puc* and *Socs36E* in ovaries following ovarian somatic cell-specific *hey* knockdown using two independent RNAi lines. **(k)** qRT-PCR analysis of *puc* and *Socs36E* in ovaries from control (*tj>w^1118^*) and ovarian somatic cell-specific *hey*-overexpressing (*tj>UAS-hey*) flies at 7, 28, and 42 days of age. Data are presented as mean ± SEM. *p* < 0.05(*) , *p* < 0.01(**), *p* < 0.001(***) , and *p* < 0.0001 (****); ns, not significant. SA-β-gal, senescence-associated β-galactosidase.

## Supplementary Methods

### Human ovarian tissue donors

Human ovarian tissues were collected from 20 participants at Women’s Hospital, Zhejiang University School of Medicine. Donors were assigned to three reproductive-stage groups using a combination of chronological age, menstrual history and serum FSH levels. The young group was defined as women aged 18-35 years with regular menstrual cycles, basal FSH <10 IU/L and anti-Müllerian hormone (AMH) ≥1.1 ng ml⁻¹ when available. The transition group was defined operationally as non-menopausal women aged 40-55 years with reduced ovarian reserve or reproductive aging features and basal FSH <25 IU/L. This group was designed to capture a late reproductive-to-menopausal transition window and was not intended to represent a strict STRAW+10-defined perimenopausal cohort. The post group included women aged 45-65 years with amenorrhoea for more than 12 months and FSH >25 IU/L.

For young donors, ovarian cortical biopsy specimens of approximately 0.5 × 0.5 × 0.2 cm were collected during clinically indicated surgery when feasible. For transition and post donors, ovarian tissues were obtained mainly from patients undergoing bilateral oophorectomy. Surgical indications included ovarian mature teratoma, benign ovarian cystic lesions, endometriosis, adenomyosis, uterine leiomyoma, tumours of uncertain malignant potential and selected early-stage non-ovarian malignancies. In all cases, sampling was restricted to macroscopically uninvolved ovarian tissue, and final pathological examination confirmed the absence of ovarian tumour involvement.

Clinical information was collected from medical records, including age, menstrual history, medical history, body mass index and reproductive hormone measurements. Serum reproductive hormones included FSH, luteinizing hormone, oestradiol and progesterone when available. AMH was recorded when available, primarily in non-menopausal participants. Some hormone measurements were missing because testing had been performed at outside hospitals or was not repeated before surgery. Donor-level clinical information and sequencing allocation are provided in Supplementary Table 1.

Participants were excluded if they had polycystic ovary syndrome or other known reproductive endocrine disorders; known chromosomal abnormalities, including Turner syndrome; use of hormone replacement therapy, cycle-regulating therapy or other steroid-containing medications within 6 months before surgery; uncontrolled endocrine, cardiovascular or autoimmune disease; abnormal liver or renal function, coagulation dysfunction or severe anaemia; previous chemotherapy or radiotherapy; severe psychiatric or cognitive disorders; or inability to provide informed consent. Participants with current early-stage non-ovarian malignancy or tumours of uncertain malignant potential were included only when ovarian tissue was macroscopically uninvolved and confirmed by final pathology to be free of tumour involvement.

Fresh ovarian tissue was placed in a pre-cooled preservation medium within 15 min after surgical removal and transported under cold-chain conditions. Visible blood clots and necrotic tissue were removed before processing. Biopsy specimens were preferentially collected from the ovarian cortex, whereas whole-ovary specimens could include both cortical and medullary regions. Tissues were processed for single-cell dissociation within 48 h after collection.

### RNA extraction and reverse transcription

Total RNA was extracted from mouse ovaries, *Drosophila* ovaries, KGN cells, and primary mouse granulosa cells using TRIzol Reagent (Thermo Fisher Scientific, Cat. No. 15596026CN). Tissue samples were homogenized using a handheld tissue grinder before RNA extraction. RNA concentration and purity were measured using a NanoDrop 2000/2000c UV-Vis spectrophotometer (Thermo Fisher Scientific), and RNA quality was assessed based on the A260/A280 ratio.

Complementary DNA (cDNA) was synthesized from total RNA using the HiScript II 1st Strand cDNA Synthesis Kit (Vazyme) according to the manufacturer’s instructions. The resulting cDNA was stored at −20°C until quantitative real-time PCR analysis.

### Quantitative real-time PCR

Quantitative real-time PCR was performed using Hieff UNICON® qPCR SYBR Green Master Mix (Yeasen Biotechnology, Cat. No. 11198ES) on a Roche LightCycler 480 II real-time PCR system equipped with a 384-well block. Reaction setup and amplification conditions were performed according to the manufacturers’ instructions.

Human gene expression was normalized to *GAPDH*, mouse gene expression was normalized to *Gapdh*, and *Drosophila* gene expression was normalized to *RpL32* (known as *rp49*). Relative gene expression was calculated using the 2^−ΔΔCt^ method. The primer sequences used are listed below.

Human qPCR primers:

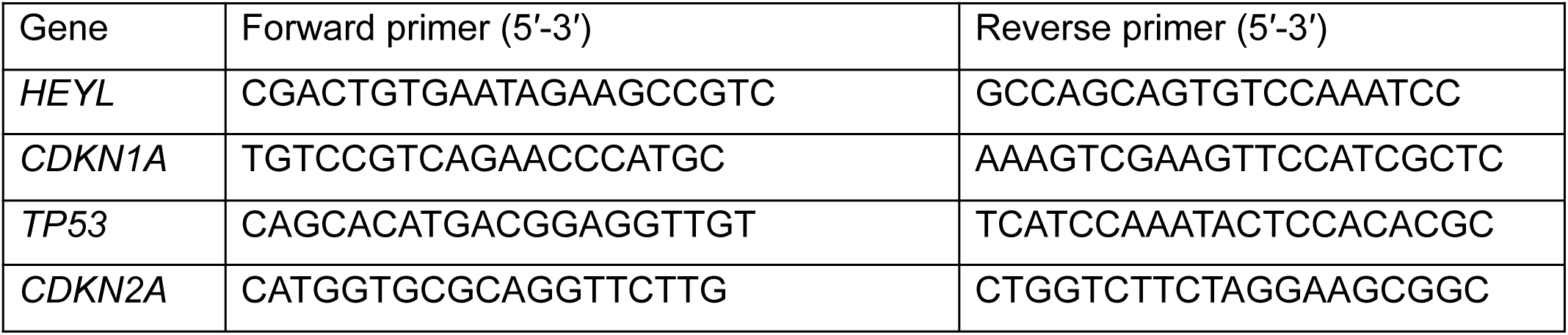
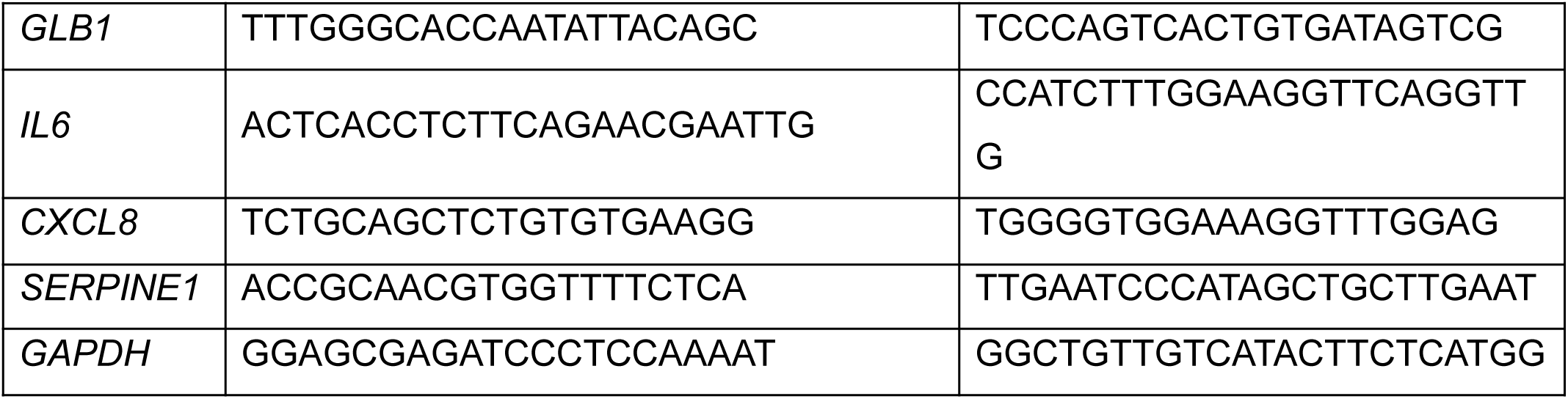

Mouse qPCR primers:

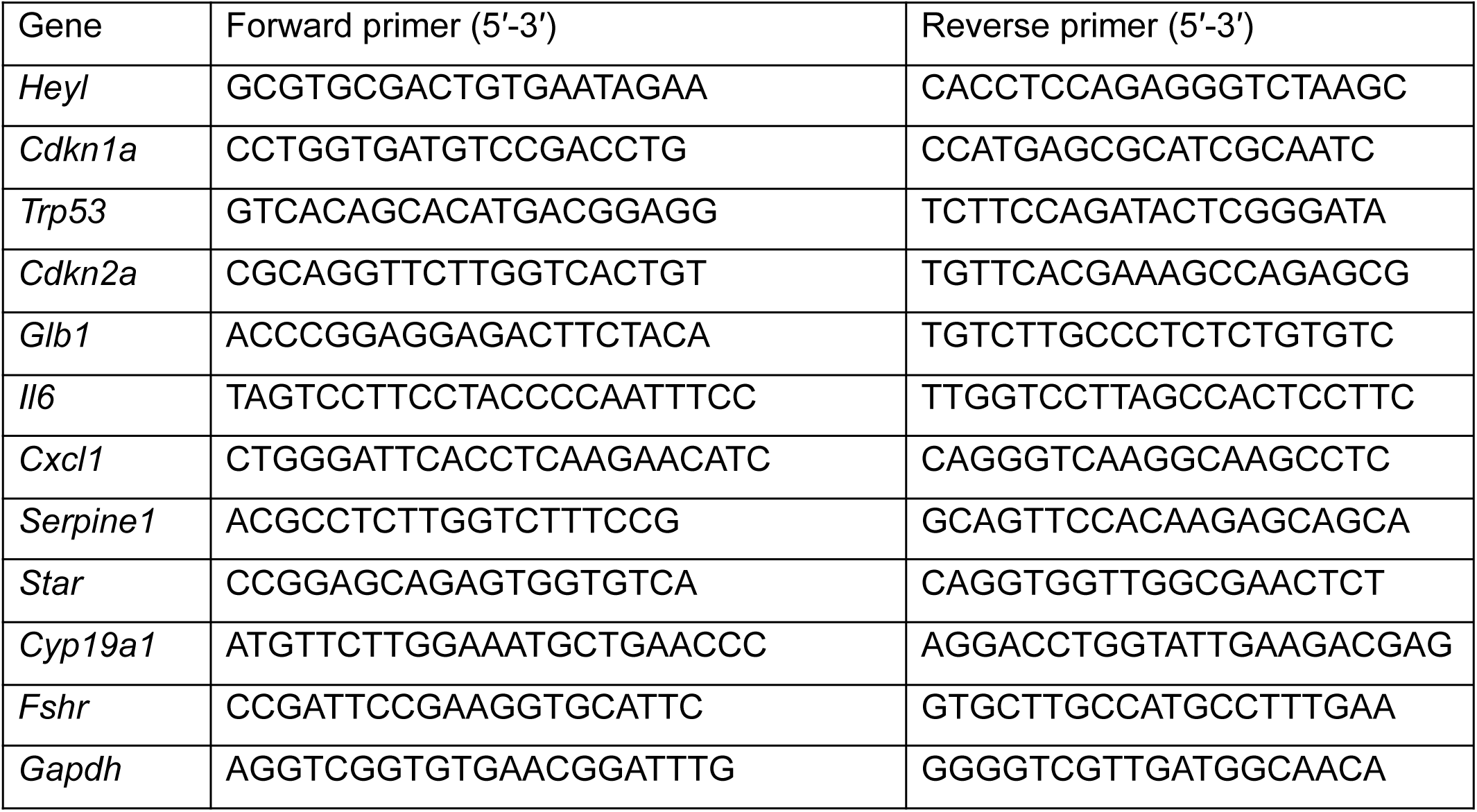

Drosophila qPCR primers:

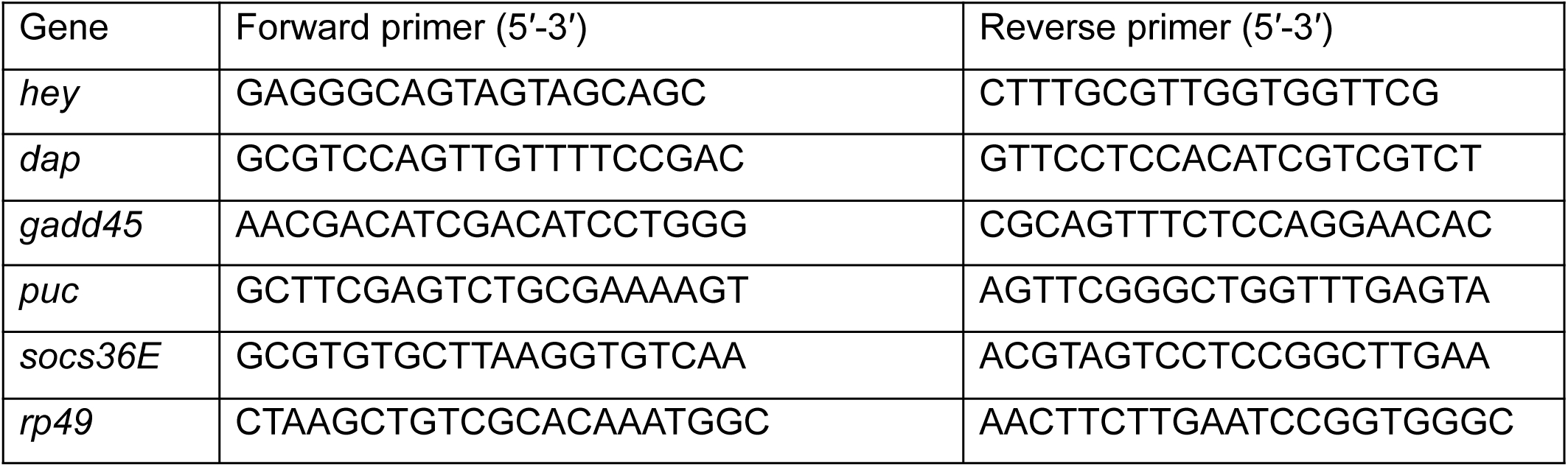

### Western blotting

KGN cells, primary mouse granulosa cells, and *Drosophila* ovaries were lysed in 4% SDS lysis buffer supplemented with a protease and phosphatase inhibitor cocktail (Beyotime, Cat. No. P1048; 50× stock solution).

Protein concentrations were determined using an Enhanced BCA Protein Assay Kit (Beyotime, Cat. No. P0010S). Equal amounts of protein were separated by SDS-PAGE and transferred onto polyvinylidene difluoride (PVDF) membranes. The membranes were blocked with 5% bovine serum albumin in Tris-buffered saline containing 0.1% Tween-20 (TBST) for 1 h at room temperature and then incubated with the indicated primary antibodies overnight at 4°C.

For KGN cells and primary mouse granulosa cells, the following primary antibodies were used: anti-HEYL/Heyl (HEYL Polyclonal Antibody; Proteintech, Cat. No. 15679-1-AP), anti-p21 (p21 Polyclonal Antibody; Proteintech, Cat. No. 28248-1-AP), and anti-p53 (p53 Antibody [DO-1]; Santa Cruz Biotechnology, Cat. No. sc-126). For *Drosophila* ovarian samples carrying the tagged *hey* transgene, tagged Hey protein was detected using an anti-FLAG antibody (DYKDDDDK Tag Polyclonal Antibody; Proteintech, Cat. No. 20543-1-AP), with α-tubulin used as the loading control (Alpha-Tubulin Monoclonal Antibody; Proteintech, Cat. No. 66031-1-Ig).

After washing, the membranes were incubated with the appropriate horseradish peroxidase-conjugated secondary antibodies for 1 h at room temperature. Protein bands were visualized using Immobilon ECL Ultra Western HRP Substrate (Millipore, Cat. No. WBULS0500), and chemiluminescence signals were captured using a ChemiScope 6000 imaging system (Clinx Science Instruments).

### KGN cell culture

The human granulosa-like tumor cell line KGN was kindly provided by the laboratory of Professor Jun Ma. Cells were cultured in Dulbecco’s modified Eagle’s medium/Nutrient Mixture F-12 (DMEM/F-12; Yuanpei Biotechnology, Shanghai, China) supplemented with 10% fetal bovine serum and 1% penicillin-streptomycin solution (100×; Yuanpei Biotechnology, Shanghai, China; Cat. No. S110JV). Cells were maintained at 37°C in a humidified incubator containing 5% CO_₂_.

When KGN cells reached approximately 70%-90% confluence, they were passaged using trypsin (Yuanpei Biotechnology, Shanghai, China; Cat. No. S310KJ).

### SA-β-galactosidase staining of cultured cells

Senescence-associated β-galactosidase staining of KGN cells and primary mouse granulosa cells was performed using a β-galactosidase staining kit (Beyotime, Cat. No. RG0039) according to the manufacturer’s instructions. Images were acquired using a Nikon ECLIPSE 80i microscope. Five randomly selected fields were analyzed for each sample, and the numbers of SA-β-gal-positive cells and total cells were counted.

The percentage of SA-β-gal-positive cells was calculated by dividing the number of SA-β-gal-positive cells by the total number of cells and multiplying by 100.

### SA-β-galactosidase staining of *Drosophila* ovaries

*Drosophila* ovaries were dissected in Schneider’s *Drosophila* Medium and stained using a β-galactosidase staining kit (Beyotime, Cat. No. RG0039) according to the manufacturer’s instructions. All samples within the same experiment were processed in parallel and imaged using identical illumination, exposure, and acquisition settings. Representative images were used for qualitative comparison of ovarian SA-β-galactosidase staining among the experimental groups.

## References

1. Dong, L., Teh, D. B. L., Kennedy, B. K. & Huang, Z. Unraveling female reproductive senescence to enhance healthy longevity. Cell Res 33, 11–29 (2023).

2. Hall, J. E. Endocrinology of the Menopause. Endocrinol Metab Clin North Am 44, 485–496 (2015).

3. López-Otín, C., Blasco, M. A., Partridge, L., Serrano, M. & Kroemer, G. Hallmarks of aging: An expanding universe. Cell 186, 243–278 (2023).

4. Lian, X. et al. Multi-omics pleiotropic association analyses reveal functionally relevant genes and druggable pathways for ovarian aging. Genome Biol. 10.1186/s13059-026-04118-7 (2026) doi:10.1186/s13059-026-04118-7.

5. Zhang, F. et al. Harnessing omics data for drug discovery and development in ovarian aging. 31, 240–268 (2025).

6. Ruth, K. S. et al. Genetic insights into biological mechanisms governing human ovarian ageing. Nature 596, 393–397 (2021).

7. Wagner, M. et al. Single-cell analysis of human ovarian cortex identifies distinct cell populations but no oogonial stem cells. Nat Commun 11, 1147 (2020).

8. Fan, X. et al. Single-cell reconstruction of follicular remodeling in the human adult ovary. Nat Commun 10, 3164 (2019).

9. Wang, S. et al. Single-Cell Transcriptomic Atlas of Primate Ovarian Aging. Cell 180, 585–600.e19 (2020).

10. Jones, A. S. K. et al. Cellular atlas of the human ovary using morphologically guided spatial transcriptomics and single-cell sequencing. Sci Adv 10, eadm7506 (2024).

11. Jin, C. et al. Molecular and genetic insights into human ovarian aging from single-nuclei multi-omics analyses. Nat Aging 5, 275–290 (2025).

12. Wu, M. et al. Spatiotemporal transcriptomic changes of human ovarian aging and the regulatory role of FOXP1. Nat Aging 4, 527–545 (2024).

13. Zhang, M. et al. Spatial Transcriptomic Characteristics of the Aging Human Ovary. Aging Cell 25, e70288 (2026).

14. Ma, Y. et al. Polygenic regression uncovers trait-relevant cellular contexts through pathway activation transformation of single-cell RNA sequencing data. Cell Genom. 3, 100383 (2023).

15. Ma, Y. et al. Integrating single-cell sequencing data with GWAS summary statistics reveals CD16+monocytes and memory CD8+T cells involved in severe COVID-19. Genome Med. 14, 16 (2022).

16. Li, J. et al. Integrating microbial GWAS and single-cell transcriptomics reveals associations between host cell populations and the gut microbiome. Nat. Microbiol. 10, 1210–1226 (2025).

17. Ma, Y. et al. Integrating polygenic signals and single-cell multiomics identifies cell-type-specific regulomes critical for immune- and aging-related diseases. *Nat*. Aging 1–20 (2025).

18. Ma, Y. Mapping genetic risk to regulatory networks at single-cell resolution. Nat. Rev. Genet. 1–2 (2026).

19. Song, L., Chen, W., Hou, J., Guo, M. & Yang, J. Spatially resolved mapping of cells associated with human complex traits. Nature 641, 932–941 (2025).

20. Mangiola, S. et al. sccomp: Robust differential composition and variability analysis for single-cell data. Proc Natl Acad Sci U S A 120, e2203828120 (2023).

21. te Velde, E. R. & Pearson, P. L. The variability of female reproductive ageing. Hum Reprod Update 8, 141–154 (2002).

22. He, L. et al. NEBULA is a fast negative binomial mixed model for differential or co-expression analysis of large-scale multi-subject single-cell data. Commun Biol 4, 629 (2021).

23. Tacutu, R. et al. Human Ageing Genomic Resources: new and updated databases. Nucleic Acids Res 46, D1083–D1090 (2018).

24. Morabito, S., Reese, F., Rahimzadeh, N., Miyoshi, E. & Swarup, V. hdWGCNA identifies co-expression networks in high-dimensional transcriptomics data. Cell Rep Methods 3, 100498 (2023).

25. Matsuda, F., Inoue, N., Manabe, N. & Ohkura, S. Follicular growth and atresia in mammalian ovaries: regulation by survival and death of granulosa cells. J Reprod Dev 58, 44–50 (2012).

26. Regan, S. L. P. et al. Granulosa Cell Apoptosis in the Ovarian Follicle-A Changing View. Front Endocrinol (Lausanne) 9, 61 (2018).

27. Lan, T. C. T. et al. Aging disrupts spatiotemporal coordination in the cycling murine ovary. Nat Aging 6, 1244–1266 (2026).

28. Gulati, G. S. et al. Single-cell transcriptional diversity is a hallmark of developmental potential. Science 367, 405–411 (2020).

29. Ferraz-de-Souza, B., Lin, L. & Achermann, J. C. Steroidogenic factor-1 (SF-1, NR5A1) and human disease. Mol Cell Endocrinol 336, 198–205 (2011).

30. Giguère, V. Transcriptional control of energy homeostasis by the estrogen-related receptors. Endocr Rev 29, 677–696 (2008).

31. Pelusi, C., Ikeda, Y., Zubair, M. & Parker, K. L. Impaired follicle development and infertility in female mice lacking steroidogenic factor 1 in ovarian granulosa cells. Biol Reprod 79, 1074–1083 (2008).

32. Eferl, R. & Wagner, E. F. AP-1: a double-edged sword in tumorigenesis. Nat Rev Cancer 3, 859–868 (2003).

33. Martínez-Zamudio, R. I. et al. AP-1 imprints a reversible transcriptional programme of senescent cells. Nat Cell Biol 22, 842–855 (2020).

34. Schep, A. N., Wu, B., Buenrostro, J. D. & Greenleaf, W. J. chromVAR: inferring transcription-factor-associated accessibility from single-cell epigenomic data. Nat. Methods 14, 975–978 (2017).

35. Bravo González-Blas, C., et al. SCENIC+: single-cell multiomic inference of enhancers and gene regulatory networks. Nat Methods 20, 1355–1367 (2023).

36. Day, F. R. et al. Genomic analyses identify hundreds of variants associated with age at menarche and support a role for puberty timing in cancer risk. Nat Genet 49, 834–841 (2017).

37. Verdiesen, R. M. G. et al. Genome-wide association study meta-analysis identifies three novel loci for circulating anti-Müllerian hormone levels in women. Hum Reprod 37, 1069–1082 (2022).

38. Rahmioglu, N. et al. The genetic basis of endometriosis and comorbidity with other pain and inflammatory conditions. Nat Genet 55, 423–436 (2023).

39. Day, F. et al. Large-scale genome-wide meta-analysis of polycystic ovary syndrome suggests shared genetic architecture for different diagnosis criteria. PLoS Genet 14, e1007813 (2018).

40. Phelan, C. M. et al. Identification of 12 new susceptibility loci for different histotypes of epithelial ovarian cancer. Nat Genet 49, 680–691 (2017).

41. Michailidou, K. et al. Association analysis identifies 65 new breast cancer risk loci. Nature 551, 92–94 (2017).

42. Komar, C. M. Peroxisome proliferator-activated receptors (PPARs) and ovarian function--implications for regulating steroidogenesis, differentiation, and tissue remodeling. Reprod Biol Endocrinol 3, 41 (2005).

43. Iso, T., Kedes, L. & Hamamori, Y. HES and HERP families: multiple effectors of the Notch signaling pathway. J Cell Physiol 194, 237–255 (2003).

44. Vanorny, D. A., Prasasya, R. D., Chalpe, A. J., Kilen, S. M. & Mayo, K. E. Notch signaling regulates ovarian follicle formation and coordinates follicular growth. Mol Endocrinol 28, 499–511 (2014).

45. Xu, J. & Gridley, T. Notch2 is required in somatic cells for breakdown of ovarian germ-cell nests and formation of primordial follicles. BMC Biol 11, 13 (2013).

46. Kim, J. et al. Peroxisome proliferator-activated receptor gamma is a target of progesterone regulation in the preovulatory follicles and controls ovulation in mice. Mol Cell Biol 28, 1770–1782 (2008).

47. Liu, Z., Castrillon, D. H., Zhou, W. & Richards, J. S. FOXO1/3 depletion in granulosa cells alters follicle growth, death and regulation of pituitary FSH. Mol Endocrinol 27, 238–252 (2013).

48. Stolk, L. et al. Meta-analyses identify 13 loci associated with age at menopause and highlight DNA repair and immune pathways. Nat Genet 44, 260–268 (2012).

49. Hao, Y. et al. Integrated analysis of multimodal single-cell data. Cell 184, 3573–3587.e29 (2021).

50. Korsunsky, I. et al. Fast, sensitive and accurate integration of single-cell data with Harmony. Nat Methods 16, 1289–1296 (2019).

51. Stuart, T., Srivastava, A., Madad, S., Lareau, C. A. & Satija, R. Single-cell chromatin state analysis with Signac. Nat Methods 18, 1333–1341 (2021).

52. Yu, G., Wang, L.-G., Han, Y. & He, Q.-Y. clusterProfiler: an R package for comparing biological themes among gene clusters. OMICS 16, 284–287 (2012).

53. Jin, S. et al. Inference and analysis of cell-cell communication using CellChat. Nat Commun 12, 1088 (2021).

54. Yu, G., Wang, L.-G. & He, Q.-Y. ChIPseeker: an R/Bioconductor package for ChIP peak annotation, comparison and visualization. Bioinformatics 31, 2382–2383 (2015).

55. Ritchie, M. E. et al. limma powers differential expression analyses for RNA-sequencing and microarray studies. Nucleic Acids Res 43, e47 (2015).

