## Supplementary Figures for "Single-cell and spatial multi-omics mapping of human ovarian aging across young and menopausal transition"

**Supplemantary Figure 1**

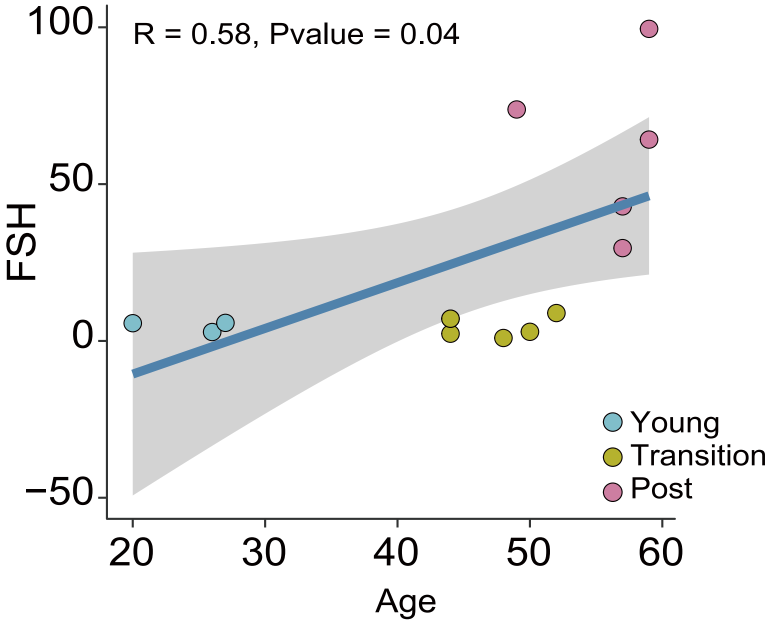


**Supplementary Figure 1: Sample FSH level correlated with age.**
Correlation between serum follicle-stimulating hormone (FSH) levels and chronological age (R = 0.58, p = 0.04).

**Supplemantary Figure 2**

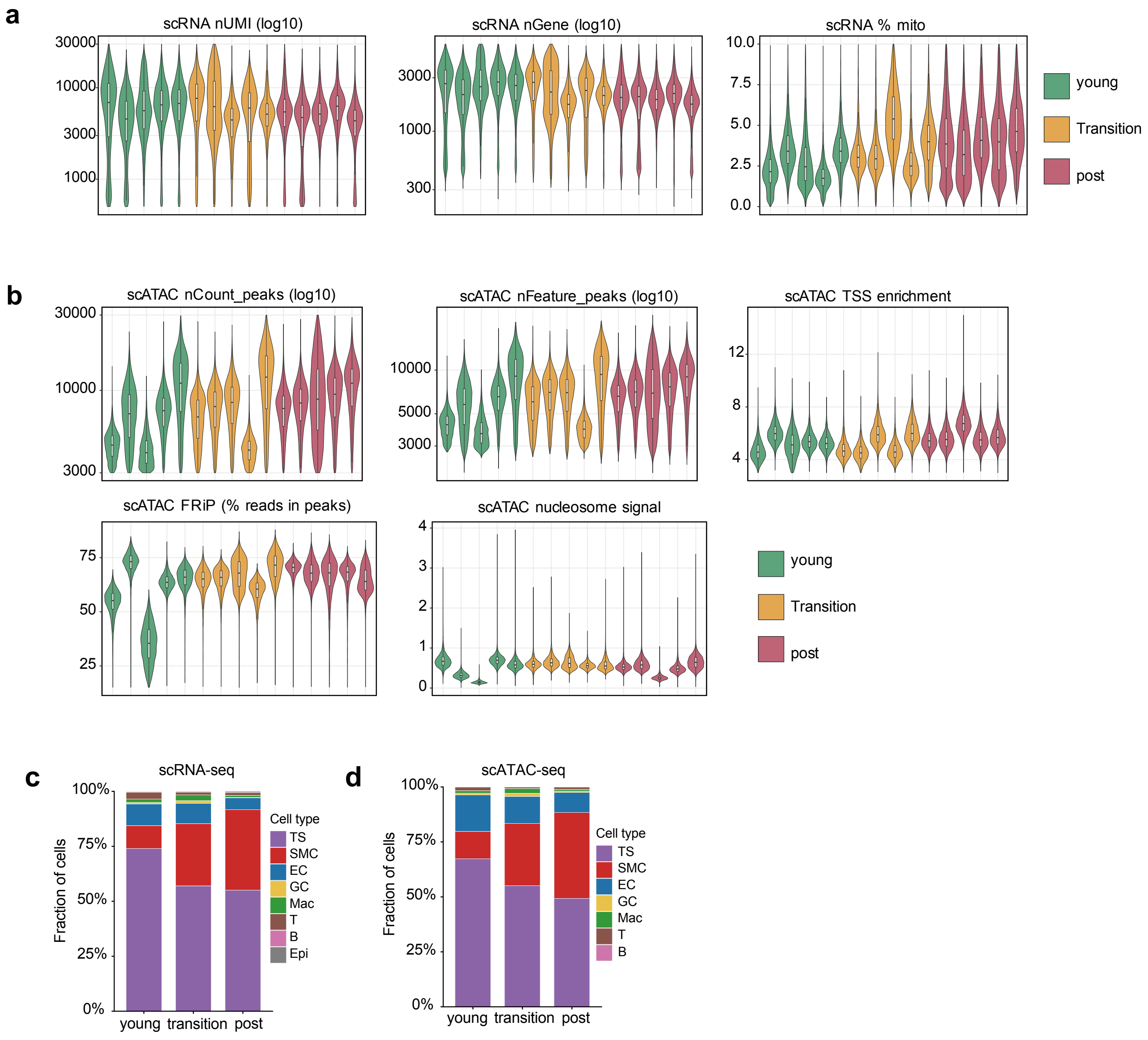


**Supplementary Figure 2: Quality control, staging, and cellular composition across age groups.** **(a,b)** Per-sample quality control metrics for scRNA-seq and scATAC-seq, ordered from young, transition to post-menopausal samples. **(c,d)** Cell-type composition across age groups for scRNA-seq **(c)** and scATAC-seq **(d).**

### Supplementary Figure 3

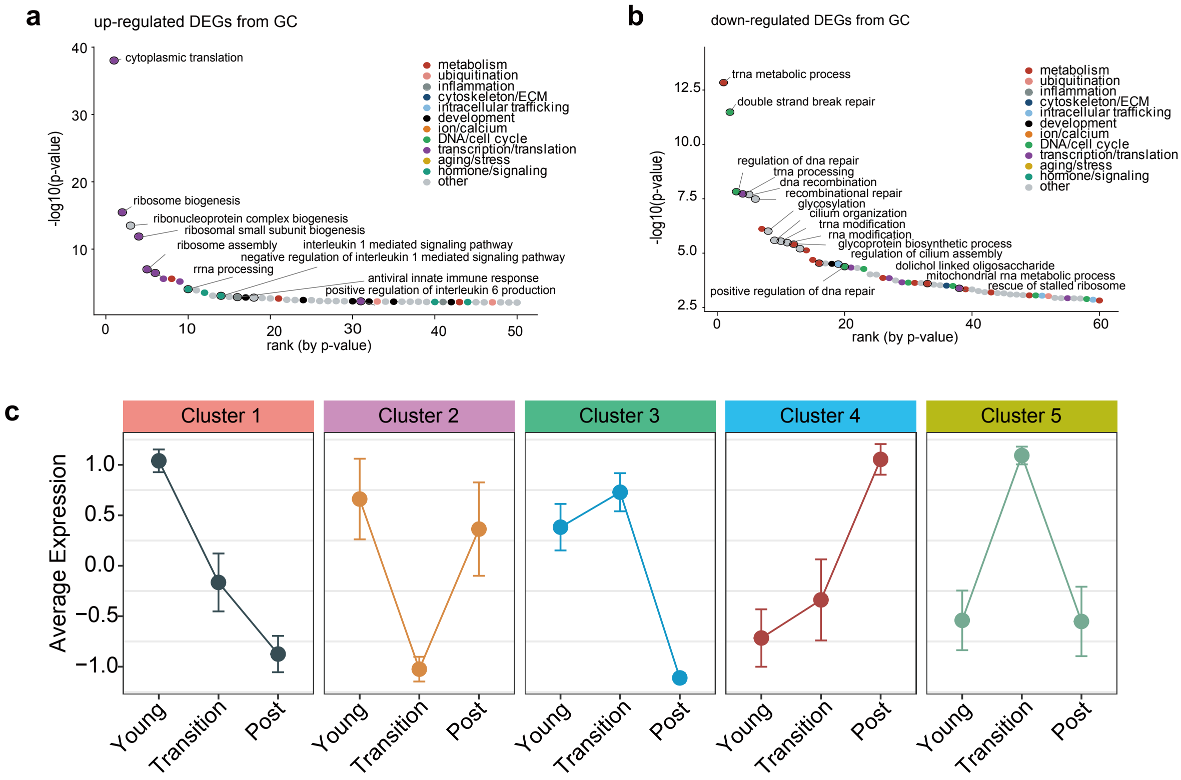


**Supplementary Figure 3: GC aging programs and their temporal dynamics.**
**(a,b)** GO enrichment of genes up-regulated **(a)** and down-regulated **(b)** with age in GC (ranked by p-value; coloured by functional category). **(c)** Average expression of each of the five aging-gene clusters across young, transition and post-menopausal groups.

**Supplementary Figure 4**

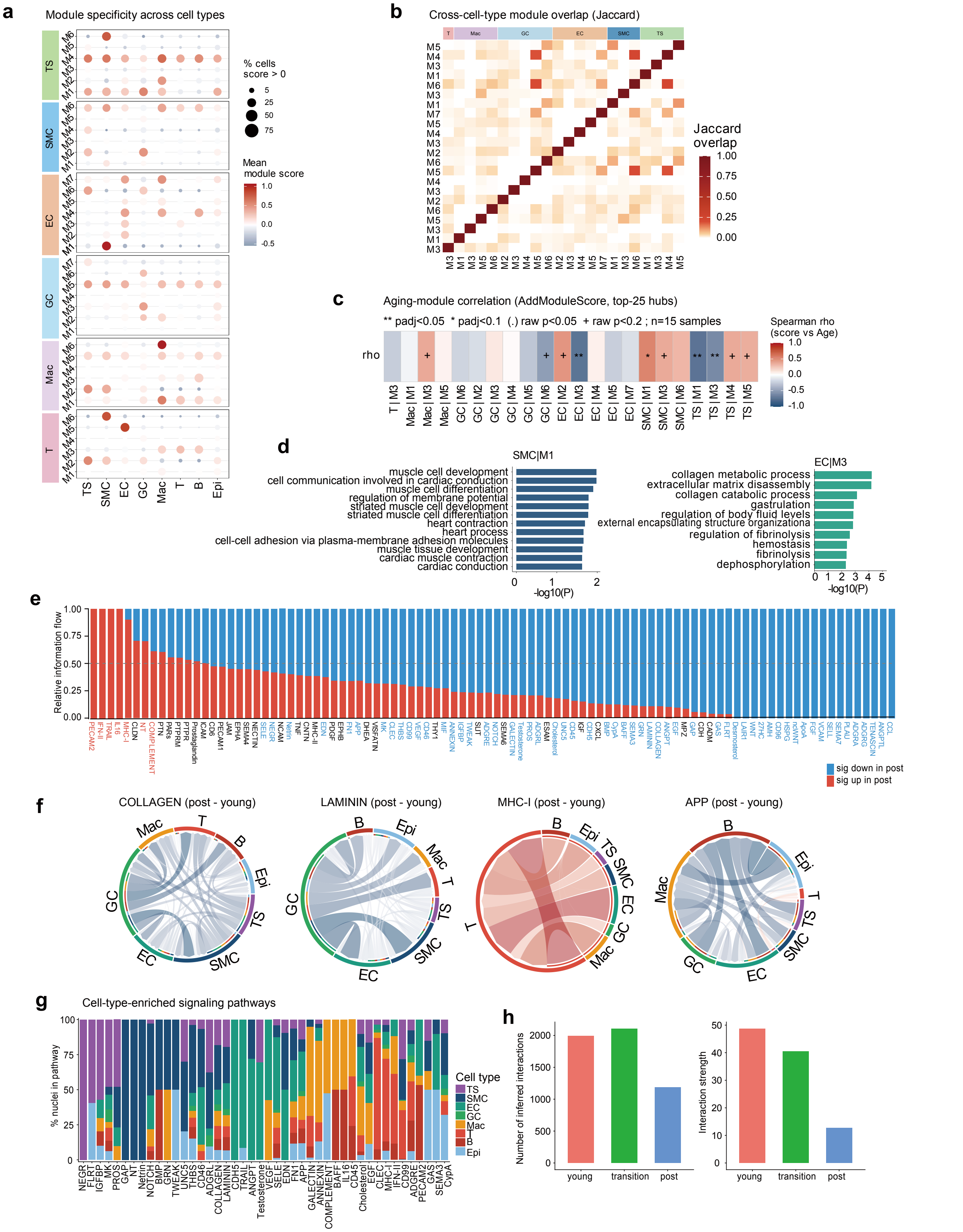


**Supplementary Fig. 4: Complete cell-type module set, module functions and age-associated cell–cell communication.**
**(a)** Module specificity across cell types for the complete set of per-cell-type hdWGCNA modules. **(b)** Cross-cell-type module overlap: Jaccard similarity between module gene sets. **(c)** Aging-module correlation: Spearman correlation between top-25 hub-gene module score (AddModuleScore) and age per cell type × module. ** $p_{adj} < 0.05$, * $p_{adj} < 0.1$, (.) raw $p < 0.05$, + raw $p < 0.2$; n = 15 samples. **(d)** GO enrichment of the smooth-muscle module SMC|M1 and the endothelial module EC|M3. **(e)** CellChat relative information flow: signaling pathways significantly up- (red) or down-regulated (blue) in post- versus young ovaries. **(f)** Differential signaling chord diagrams for COLLAGEN, LAMININ, MHC-I and APP (post v.s. young). **(g)** Cell-type contribution (% of nuclei) to each significantly enriched signaling pathway in the post-menopausal group. **(h)** Total number of inferred interactions and overall interaction strength in young, transition and post-menopausal ovaries.

### Supplementary Figure 5

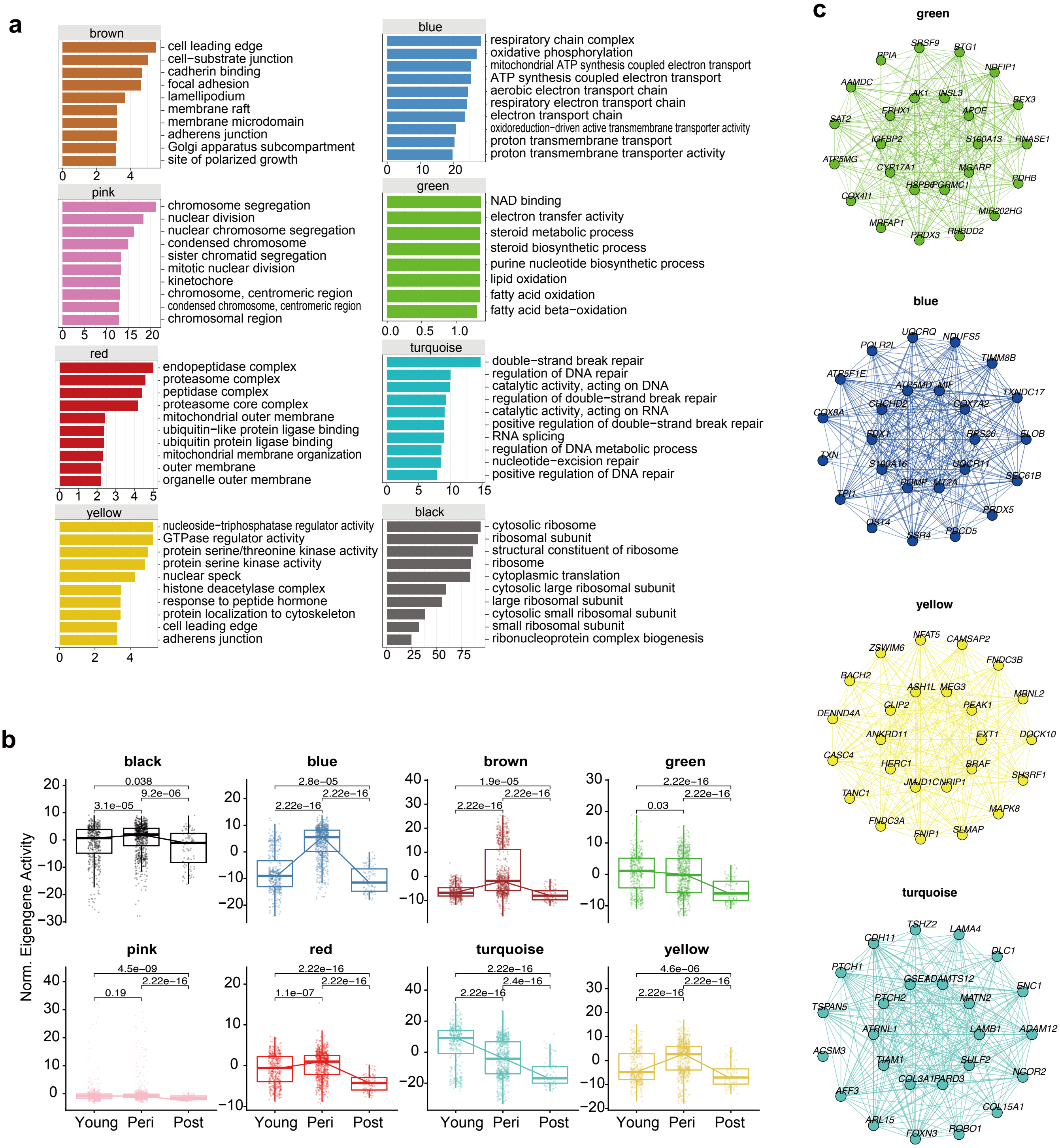


**Supplementary Fig. 5: GC co-expression module function, aging dynamics and hub-gene networks. (a)** Gene Ontology enrichment for each GC hdWGCNA module. **(b)** Module eigengene activity by age group (young/transition/post) for each module. **(c)** Hub-gene co-expression networks for representative modules.

### Supplementary Figure 6

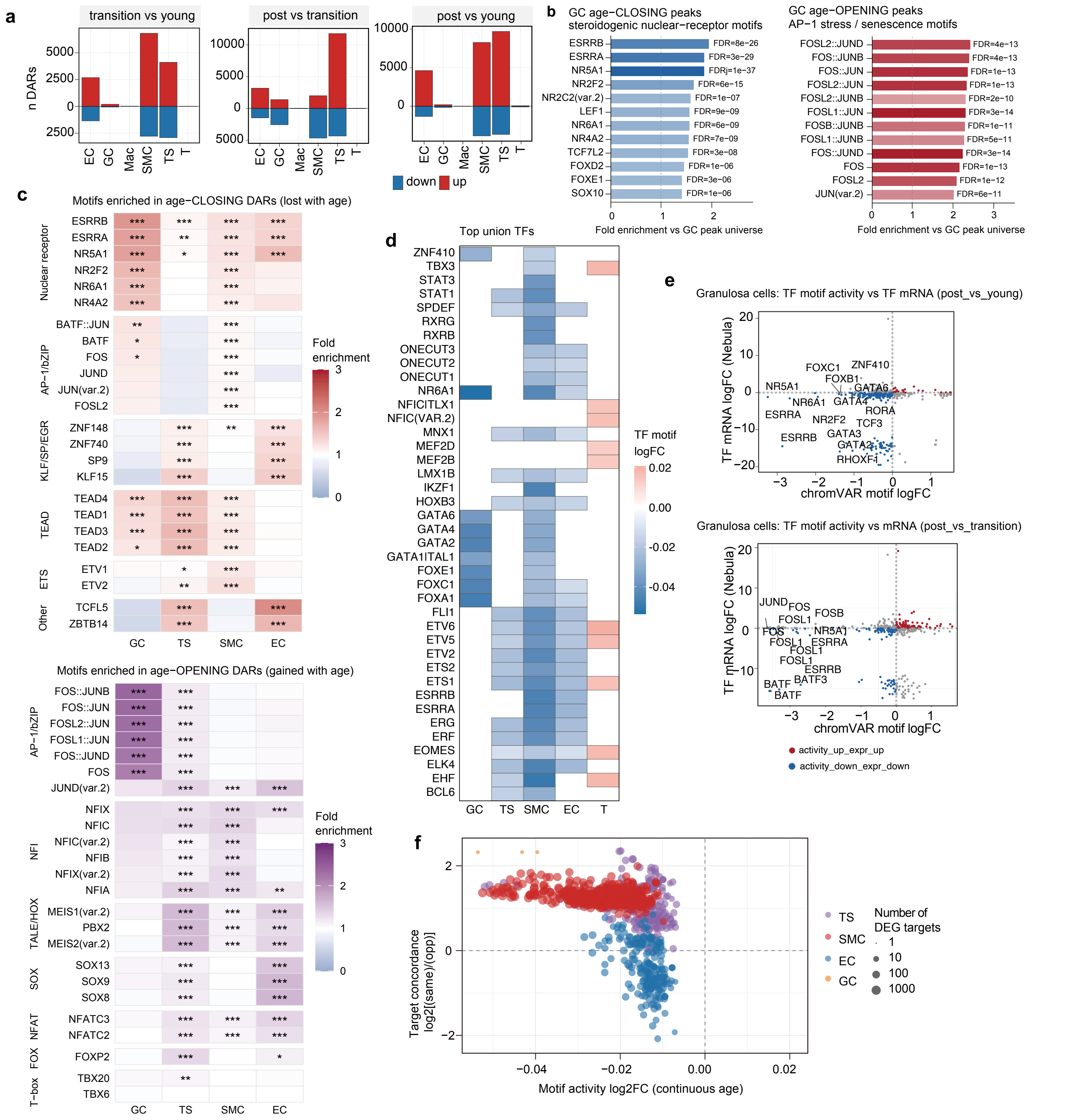

**Supplementary Fig. 6: Genome-wide DAR characterization and transcription-factor-activity dynamics during ovarian aging.** (a) Number of differentially accessible regions (DARs) per cell type for each pairwise age comparison (transition vs young, post vs transition, post vs young; logistic-regression test, Bonferroni-adjusted $p < 0.05$, $|\log2FC| > 0.1$). Bars above/below zero denote gained (up, red) and lost (down, blue) accessibility. (b) Transcription-factor motifs enriched in GC age-associated DARs (one-sided Fisher test against the granulosa tested-peak universe; JASPAR2020 CORE motifs). Left, motifs enriched among peaks that lose accessibility with age: nuclear-receptor motifs (ESRRB, ESRRA, NR5A1, NR2F2, NR4A2); right, motifs enriched among peaks that gain accessibility: AP-1/bZIP motifs (FOS, JUN, FOSL and JUNB dimers). (c) Cell-type-resolved motif enrichment in age-associated DARs, computed with the same uniform one-sided Fisher test against each cell type's own tested-peak background (motifmatchr; JASPAR2020 CORE). Top, age-closing peaks; bottom, age-opening peaks; rows grouped by TF family. asterisks = adjusted p (* < 0.05, ** < 0.01, *** < 0.001). (d) Heatmap of motif-activity change with continuous age (chromVAR deviation regressed on age) for the top union of differentially active transcription factors (n = 40) across cell types. Color denotes the age coefficient of TF motif activity. (e) Granulosa-cell TF motif activity (chromVAR log2(fold-change)) versus TF mRNA log2(fold-change) (NEBULA) for post vs young (top) and post vs transition (bottom). Points are colored by fold-change directions. (f) Directional concordance between TF motif activity and target-gene expression. For each TF, the log2 ratio of its differentially expressed (DEG) targets changing in the same versus the opposite direction to its motif-activity change is plotted against the motif-activity log2(fold-change) with continuous age.

### Supplementary Figure 7

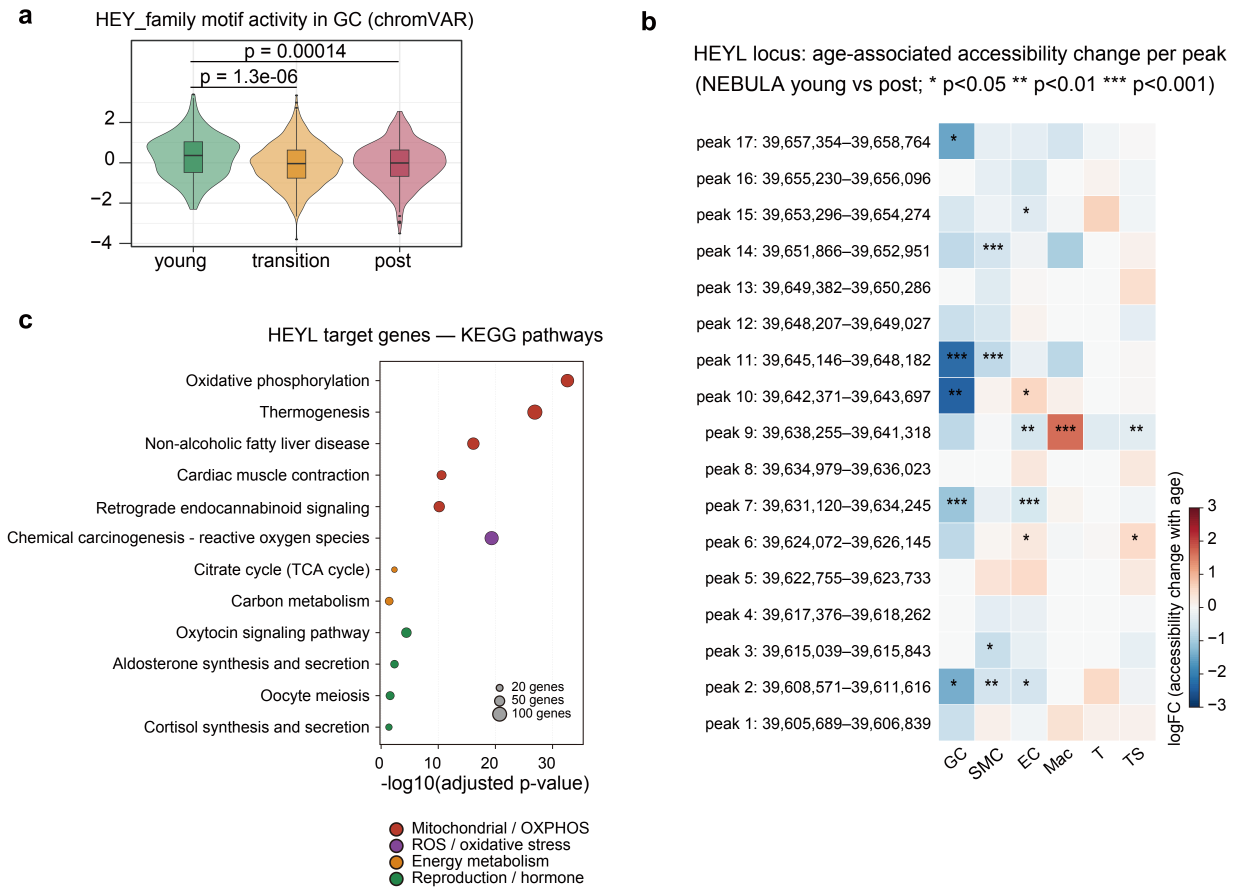


**Supplementary Fig. 7: *HEYL*-locus accessibility, transcription-factor activity and target pathways.** (a) *HEY*-family transcription-factor motif activity (chromVAR deviation) in GC across young, transition and post-menopausal groups. Violin plots show per-cell motif z-scores with pairwise Wilcoxon rank-sum p values. (b) Heatmap of the age-associated accessibility change (NEBULA log2(fold-change), young vs post) for all 17 *HEYL*-locus peaks (rows, by genomic position) across cell types (columns). *$p < 0.05$, **$p < 0.01$, ***$p < 0.001$. (c) KEGG pathway enrichment of HEYL SCENIC+ GRNBoost target genes (aging- and reproduction-related pathways). Dot color denotes pathway category and dot size the number of contributing genes.

### Supplementary Figure 8

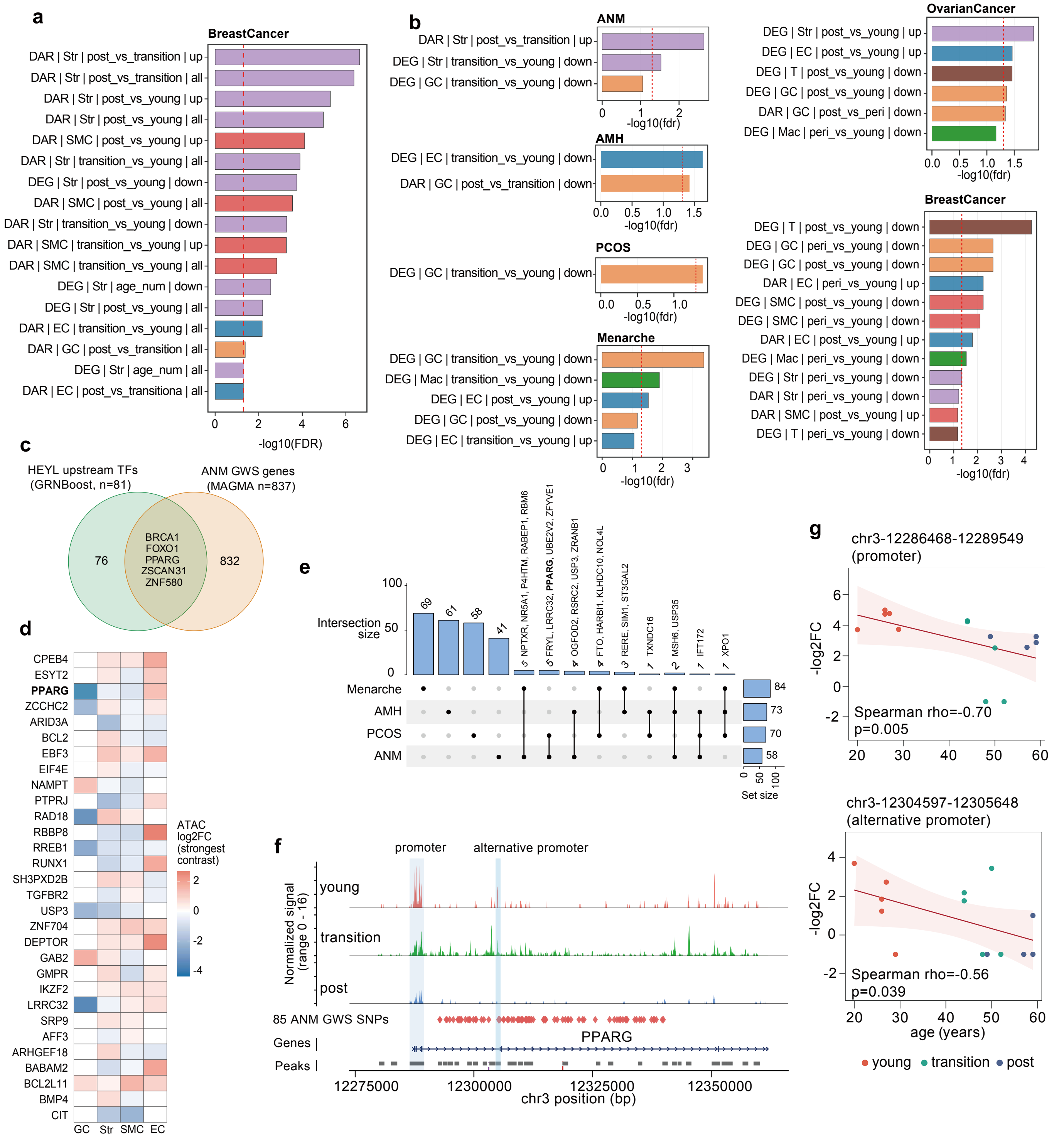


**Supplementary Fig. 8: Ovarian trait gene enrichment in age-remodeled chromatin, the *HEYL* regulatory circuit and the PPARG locus.**
**(a)** Enrichment of ovarian trait (Breast cancer) genes in age-associated DAR/DEG gene sets across cell types, age contrasts and directions. Extension of Fig. 5a. **(b)** The same enrichment using curated GWAS Catalog trait genes as an independent gene-set source (ANM, AMH, PCOS, menarche, ovarian cancer and breast cancer). **(c)** Venn diagram of *HEYL* upstream transcription factors (GRNBoost, n = 81) and ANM genome-wide-significant genes (MAGMA, Ruth2021 n = 837; Bonferroni $p < 0.05$). **(d)** Heatmap of ATAC log2(fold-change) (strongest age contrast) for the top 30 ANM-associated age-DAR genes across ovarian cell types. **(e)** UpSet plot of shared genes among the ovarian trait (menarche, AMH, PCOS, ANM) gene sets; intersection sizes (top) and per-trait set sizes (right). **(f)** ATAC coverage across the *PPARG* locus (chr3) in young, transition and post-menopausal ovaries, with the 85 ANM genome-wide-significant SNPs and the 2 aging-associated peaks and their annotations indicated. **(g)** Sample-level GC accessibility of the 2 aging-associated peaks, PPARG promoter peak (chr3:12,286,468–12,289,549; Spearman $\rho = -0.70$, $p = 0.005$) and alternative promoter peak (chr3:12,304,597–12,305,648; $\rho = -0.56$, $p = 0.039$) versus donor age.

### Supplementary Figure 9

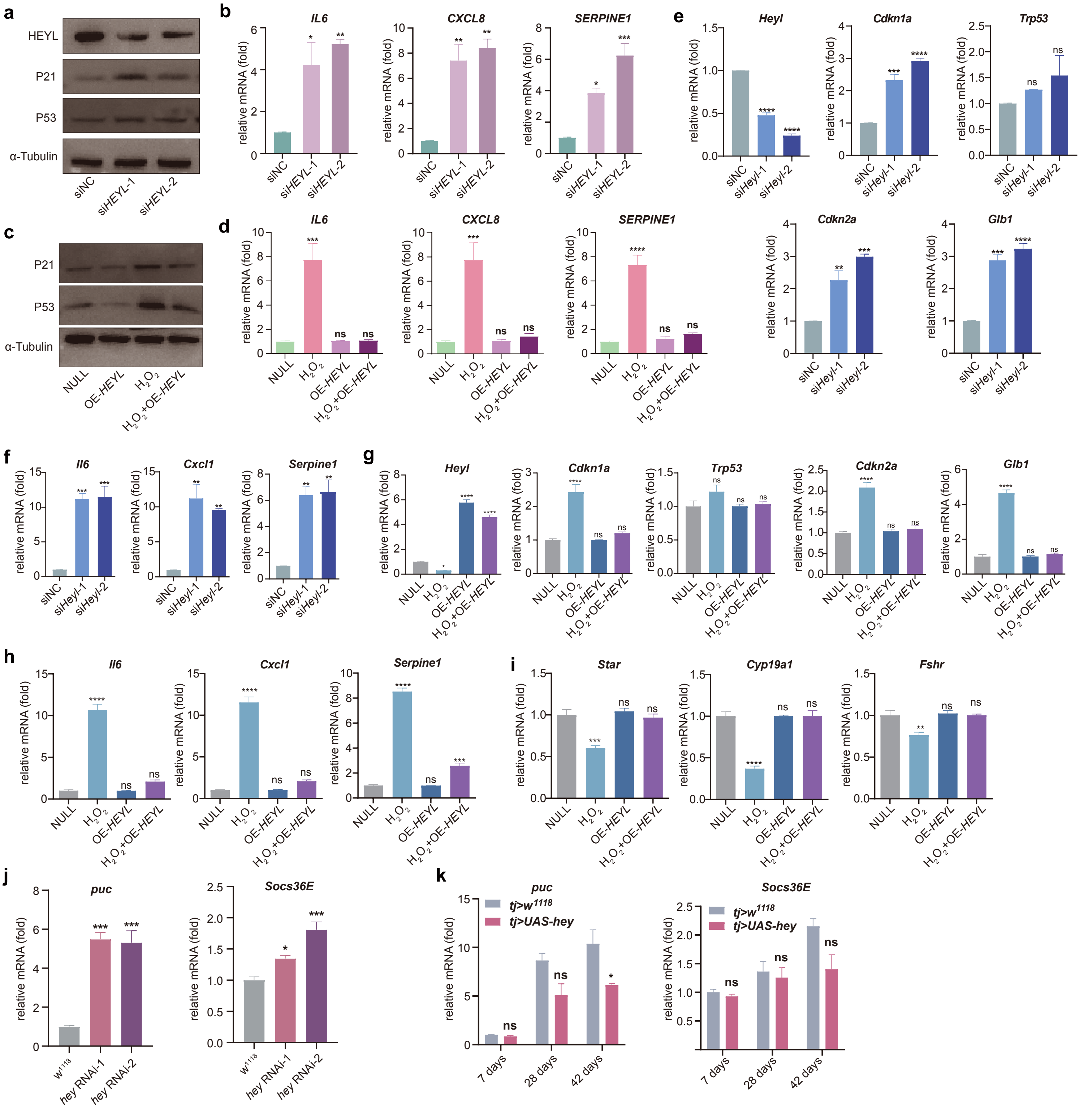


**Supplementary Figure 9: HEYL and Hey modulate senescence-associated gene expression in granulosa cells and Drosophila ovaries**

**(a)** Western blot analysis of HEYL, p21, and p53 in KGN cells transfected with control siRNA (siNC) or two independent HEYL-targeting siRNAs (si*HEYL*-1 and si*HEYL*-2). α-Tubulin served as the loading control. **(b)** qRT–PCR analysis of the SASP-related genes *IL6*, *CXCL8*, and *SERPINE1* in KGN cells following *HEYL* knockdown. **(c)** Western blot analysis of p21 and p53 in KGN cells with or without *HEYL* overexpression and $H_2O_2$ treatment. α-Tubulin served as the loading control. **(d)** qRT–PCR analysis of *IL6*, *CXCL8*, and *SERPINE1* in KGN cells under the same treatment conditions. **(e, f)** qRT–PCR analysis of *Heyl* and the senescence-associated genes *Cdkn1a*, *Trp53*, *Cdkn2a*, and *Glb1* **(e)**, and the SASP-related genes *Il6*, *Cxcl1*, and *Serpine1* **(f)** in primary mouse granulosa cells following *Heyl* knockdown using two independent siRNAs. **(g–i)** qRT–PCR analysis of *Heyl*, *Cdkn1a*, *Trp53*, *Cdkn2a*, and *Glb1* **(g)**, the SASP-related genes *Il6*, *Cxcl1*, and *Serpine1* **(h)**, and the granulosa cell functional genes *Star*, *Cyp19a1*, and *Fshr* **(i)** in primary mouse granulosa cells with or without *Heyl* overexpression and $H_2O_2$ treatment. **(j)** qRT–PCR analysis of the aging- and stress-associated genes *puc* and *Socs36E* in ovaries following ovarian somatic cell-specific hey knockdown using two independent RNAi lines. **(k)** qRT–PCR analysis of *puc* and *Socs36E* in ovaries from control (*tj*>$w^{1118}$) and ovarian somatic cell-specific hey-overexpressing (*tj*>*UAS-hey*) flies at 7, 28, and 42 days of age. Data are presented as mean ± SEM. $p < 0.05$(*) , $p < 0.01$(**), $p < 0.001$(***) , and $p < 0.0001$ (****); ns, not significant. SA-β-gal, senescence-associated β-galactosidase.
